# Systematic revision of the speciose sea slug genus *Doto* (Heterobranchia: Nudibranchia) - from the Mediterranean to South America

**DOI:** 10.1101/2025.05.14.653969

**Authors:** Diego Vázquez-Alcaide, Xavier Salvador, Gonzalo Giribet, Yuri Hooker, Michael Schrödl, Juan Moles

**Affiliations:** Departament de Biologia Evolutiva, Ecologia i Ciències Ambientals, Facultat de Biologia, Barcelona, Catalonia, Spain; EMBIMOS, Department of Physical & Technological Oceanography (CSIC), Institute of Marine Sciences (ICM), Barcelona, Catalonia, Spain; Museum of Comparative Zoology, Department of Organismic and Evolutionary Biology, Harvard University, Cambridge, Boston, USA; SNSB-Bavarian State Collection of Zoology, Section Mollusca, Münchhausenstrasse Munich, Germany; Biozentrum Ludwig Maximilians University and GeoBio-Center LMU Munich, Germany; Institut de Recerca de la Biodiversitat (IRBio), Universitat de Barcelona (UB), Barcelona, Catalonia, Spain; Universidad Peruana Cayetano Heredia, Laboratorio de Biología Marina, Facultat de Ciencias y Filosofía, Lima, Peru

**Keywords:** Gastropoda, integrative taxonomy, morphological variability, new species, phylogeny, synonymy

## Abstract

The genus *Doto* Oken, 1815 is, taxonomically, one of the most complex genera of nudibranchs due to the cryptic nature of its species, their small body size, and the homogeneity among their internal and external features. Here, an extensive molecular analysis of Mediterranean, Northeastern Atlantic, and South American specimens sheds light on the species-level taxonomy of the group. Our multilocus analyses include 171 specimens, 59 newly sequenced, corresponding to 20 species. Ten species are included in a molecular phylogeny for the first time, two being new species. Here, we provided detailed morphological and ecological descriptions of three Atlantic species, 11 Mediterranean, and one from the Pacific, complemented by live photographs evidencing their chromatic variation. The phylogenetic and species delimitation analyses suggest that coloration and external morphology often seem unreliable to differentiate among species. Molecular evidence corroborates and expands the geographical distribution of *Doto* species, some of which have never been included in molecular studies. The Mediterranean species currently recognized as *D. coronata* (Gmelin, 1791) and *D. dunnei* Lemche, 1976 correspond to *D. millbayana* Lemche, 1976. Consequently, we suggest the junior synonymy with *D. dunnei* **syn. nov.** Nevertheless, we found evidence for a restricted distribution of *D. coronata* in the western Mediterranean, coexisting in sympatry with *D. cavernicola* **sp. nov.** An additional new species from the Chilean Patagonia is also described as *D. vrenifossorum* **sp. nov.** Considering the present phylogenetic scenario, there is a highlighted need for further new morphological and genetic evidence, expanding the number of taxa and species to further unravel the taxonomy of *Doto*.

This article is registered in ZooBank under https://zoobank.org/References/99a6148d-a93c-4d2d-947e-26a85329a57f

## Introduction

Sea slugs of the order Nudibranchia are molluscs with a cosmopolitan distribution and a wide ecological and morphological diversity (e.g., Moles & Giribet, 2021). Currently, thanks to key systematic studies, the relationships between many nudibranch families appear well resolved (Wägele & Willan, 2000; Pola & Gosliner, 2010; Johnson & Gosliner, 2012; Carmona et al., 2013; Moles et al., 2023; Paz-Sedano et al., 2024) and new species are regularly being discovered (e.g., Korshunova et al., 2020; Schächinger et al., 2022; Cunha et al., 2023). The family Dotidae comprises four genera: *Caecinella* Bergh, 1870; *Doto* Oken, 1815; *Kabeiro* Shipman & Gosliner, 2015; and *Miesea* Er. Marcus, 1961. The genera *Caecinella* and *Miesea* are monotypic and their taxonomic status is considered doubtful (Odhner, 1936; Thiele, 1931). *Doto*, with a cosmopolitan distribution and 92 valid species (WoRMS, 2024), is the most diverse genus of the family. The family belongs to Dendronotoidea, within Cladobranchia (Wägele & Willan, 2000; Bouchet & Rocroi, 2005; Pola & Gosliner, 2010), with members typically displaying a rhinophoral sheath, a cuticle covering the stomach, and tentacular expansions on the oral veil (Wägele & Willan, 2000). However, unlike the rest of Dendronotoidea, Dotidae lacks the latter two characters (Wägele & Willan, 2000).

*Doto* species exhibit a distinctive morphology, featuring a small limaciform body rarely surpassing one centimetre in length (Thompson et al., 1990; Shipman & Gosliner, 2015). They possess paired dorsolateral appendages along the body, which lack cnidosacs (Martin et al., 2009) and, as such, cannot be referred to as cerata *sensu stricto*. Instead, these appendages have tubercles organized in circles known as crowns, stacked one above the other (Schmekel & Portmann, 1982; Thompson et al., 1990). On the anterior side of the body, they bear smooth finger-shaped rhinophores, surrounded at the base by a cup-shaped sheath and a smooth oral veil, which usually has two rounded lateral extensions (Moles et al., 2016). The radula is uniseriate and narrow (Thompson et al., 1990; Schmekel & Portmann, 1982). Due to significant intraspecific variation and interspecific similarity, the radula is generally not considered when differentiating between *Doto* species (Schmekel & Kress, 1977; Schmekel & Portmann, 1982; Ortea & Bouchet, 1989; Thompson et al., 1990). This species is usually defined based on external characters, such as colouration, number and shape of dorsolateral appendages, and morphology of the rhinophore sheath (Ortea & Urgorri, 1978; Lemche, 1976; Ortea & Pérez, 1982; Ortea & Bouchet, 1989). Instead, the reproductive system and their prey can also be determinants for species identification. Some species specialise in certain taxa of thecate and athecate hydrozoans (Picton & Brown, 1981; Morrow et al., 1992; Martinsson et al., 2021). They do not prey directly on the polyps of hydroids but cut the perisarc beneath the hydroid stalks and extract the fluid from the coenosarc (Thompson & Brown, 1984). The reproductive system provides systematic and phylogenetic information on *Doto* species (Shipman & Gosliner, 2015; Moles et al., 2016), as in other genera of nudibranchs (Gosliner, 1994; Wägele & Willian, 2000; DaCosta et al., 2007).

*Doto coronata* (Gemelin, 1791) is the type species of the genus, and, subsequently, through observations of specimens collected at different localities in the North Atlantic, it was determined a complex of species (Lemche, 1976). Based on the morphology and specific prey preference of hydroids, five species were described within the *D. coronata* complex: *D. dunnei* Lemche, 1976, *D. eireana* Lemche, 1976, *D. koenneckeri* Lemche, 1976, *D. millbayana* Lemche, 1976, and *D. tuberculata* Lemche, 1976. Over time, Morrow et al. (1992) first sequenced North Atlantic specimens, describing two additional species within this complex, namely *D. sarsiae* Morrow, Thorpe & Picton, 1992 and *D. hydrallmaniae* Morrow, Thorpe & Picton, 1992. Later, most of the species distinguished by Lemche (1976) and other *Doto* species from the Atlantic, Mediterranean, Indo-Pacific, and Antarctica were validated (Pola & Gosliner, 2010; Shipman & Gosliner, 2015; Moles et al., 2016; Martinsson et al., 2021). However, no significant evidence of molecular divergence between *D. millbayana* and *D. dunnei* was found (Shipman & Gosliner, 2015; Martinsson et al., 2021). Based on their molecular results, Shipman & Gosliner (2015) suggested the latter two species as synonyms. However, these authors proposed that further studies incorporating more specimens and genes from both morphotypes were needed to verify their identity. Much recently, Martinsson et al. (2021) increased the number of North Atlantic specimens of *D. dunnei* and *D. millbayana,* and their genetic results showed no divergence for both species, supporting the synonymy suggested by Shipman & Gosliner (2015). Nevertheless, they kept both species distinct because of differences in radular characters, diet, pseudobranch shape, and body colouration (Martinsson et al., 2021).

While *D. coronata* is considered widespread in the Mediterranean, sequenced material from the region did not cluster with any *D. coronata* specimens sequenced from the type locality in the Netherlands, nor the North Sea and the Northwestern Atlantic (Shipman & Gosliner, 2015; Moles et al., 2016). These molecular results suggest the Mediterranean specimens identified as *D. coronata* belong to another species, but a morphological assessment and a wider molecular framework are required to assess it. Records of ‘*D. coronata*’ in the Mediterranean are mainly taken from specimens living in *Posidonia* meadows (Sammut & Perrone, 1998; GROC, 2024; OPK-Opistobranquis, 2024). These Mediterranean specimens possess morphological differences from the original description of the species (Moles et al., 2016), such as the smoother cerata without marked tubercles (except for the apical one) and an irregular distribution of maroon markings between the tubercles.

Although *Doto* encompasses a cosmopolitan distribution (GBIF, 2024), phylogenetic studies have mainly focused on specimens collected in the Northeast Atlantic (Shipman & Gosliner, 2015; Martinsson et al., 2021). In this region, 20 species have been documented (Calado et al., 2003; Cervera et al., 2004; Picton & Morrow, 2023), 14 of which are considered autochthonous: *D. arteoi* Ortea, 1978, *D. cuspidata* Alder & Hancock, 1862, *D. eireana*, *D. hydrallmaniae*, *D. hystrix* Picton & G. H. Brown, 1981, *D. lemchei* Ortea & Urgorri, 1978, *D. maculata* (Montagu, 1804), *D. millbayana*, *D. oblicua* Ortea & Urgorri, 1978, *D. onusta* Hesse, 1872, *D. pinnatifida* (Montagu, 1804), *D. sarsiae*, *D. tuberculata*, and *D. verdicioi* Ortea & Urgorri, 1978. Out of the 20 Northeast Atlantic species, molecular data is available for 13, enabling their inclusion in phylogenetic studies. In contrast, the Coral Triangle is suggested as the most diverse area for Dotidae (Shipman & Gosliner, 2015), with 24 species documented, albeit many being undescribed (Ortea, 1982; Gosliner et al., 2008). Of these, only two (*D. greenamyeri* Shipman & Gosliner, 2015 and *D. ussi* Ortea, 1982) are molecularly verified (Shipman & Gosliner, 2015). The scant molecular data for the tropical Indo-Pacific is also evident in the Mediterranean region. In the Mediterranean, up to 15 species have been documented (Cervera et al., 2004; GROC, 2024; OPK-Opistobranquis, 2024), but only four of these have been molecularly verified (*D. dunnei*, *D. floridicola*, *D. koenneckeri*, and *D. paulinae* Trinchese, 1881) (Moles et al., 2016). The unevenness in molecular data across different regions highlights the importance of increasing the taxon sampling in poorly studied areas, such as the Mediterranean. This is crucial to achieve a global, comprehensive understanding of the diversity and evolutionary history of the genus *Doto*.

Many studies have highlighted the importance of incorporating morphology along with molecular analyses to help understand evolutionary relationships within heterobranch taxa (e.g., Fernández-Vilert et al., 2021; Tibiriçá et al., 2023; Paz-Sedano et al., 2024). In this study, our main objective is to provide molecular and morphoanatomical evidence to resolve the problematic taxonomy of the genus *Doto*. We increase the taxon sampling and geographical breadth of *Doto* species from previously published studies, including samples from the Northwest (NW) Iberian Peninsula, the Canary Islands, Antarctica, the Southwest Pacific (fiords of Patagonia, Chile), and the Eastern Mediterranean. Due to our affiliations, sampling efforts are focused on the Mediterranean, as we intended to characterize the identity of species better both morphologically and with molecular data to place them in a larger phylogenetic context.

## Methods

### Taxon Sampling

Most *Doto* specimens were collected from localities along the Western Mediterranean Coast and NE Atlantic, including the Canary Islands, Galicia and the Catalan coast (Fig. 1). Additional specimens were collected from various localities in Perú, Chile, and the South Shetland Islands in Antarctica (Fig. 2). Specimens were collected by SCUBA diving and snorkelling at a depth of 0–30 m. Images of living specimens were taken with a Nikon D90 or D7200 coupled with a 60– and 105-mm macro lens or a 10x wet lens. After collection, the specimens were fixed in 96% EtOH for molecular analysis. All specimens have been deposited at the Museum of Comparative Zoology, Harvard University (MCZ), the Animal Biodiversity Resource Centre, University of Barcelona (CRBA) and the Bavarian State Collection of Zoology of Munich (ZSM). Collecting permits were issued by the Catalan (SF/0589/2018, DG051201-333/2022) and Canary Governments (SGBTM/BDM/AUTSPP/13/2023), in addition to the ones already provided to the MCZ and the ZSM. Life cycle data were obtained from the GROC database (https://opistobranquis.org/en/home), comprising 2,109 records of the Mediterranean species under study.

**Fig. 1.**
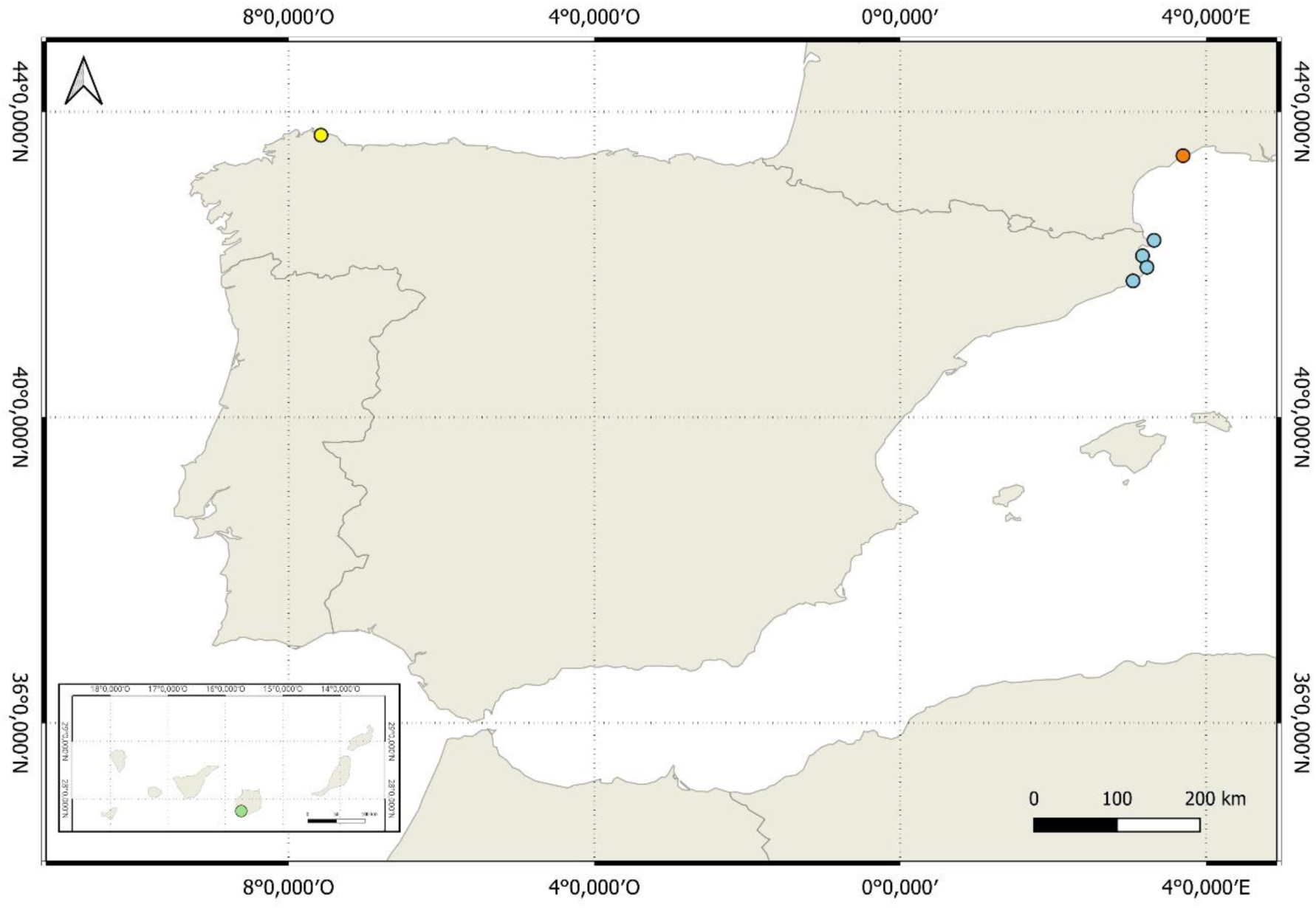
Distribution map from the Eastern Mediterranean and North Atlantic of the *Doto* species sequenced in this study. Circles represent sampling stations color-coded by region: France (orange); Catalonia, Spain (blue); Galicia, Spain (yellow); Gran Canaria, Canary Islands (green). Map generated using QGIS v.3.36.0

**Fig. 2.**
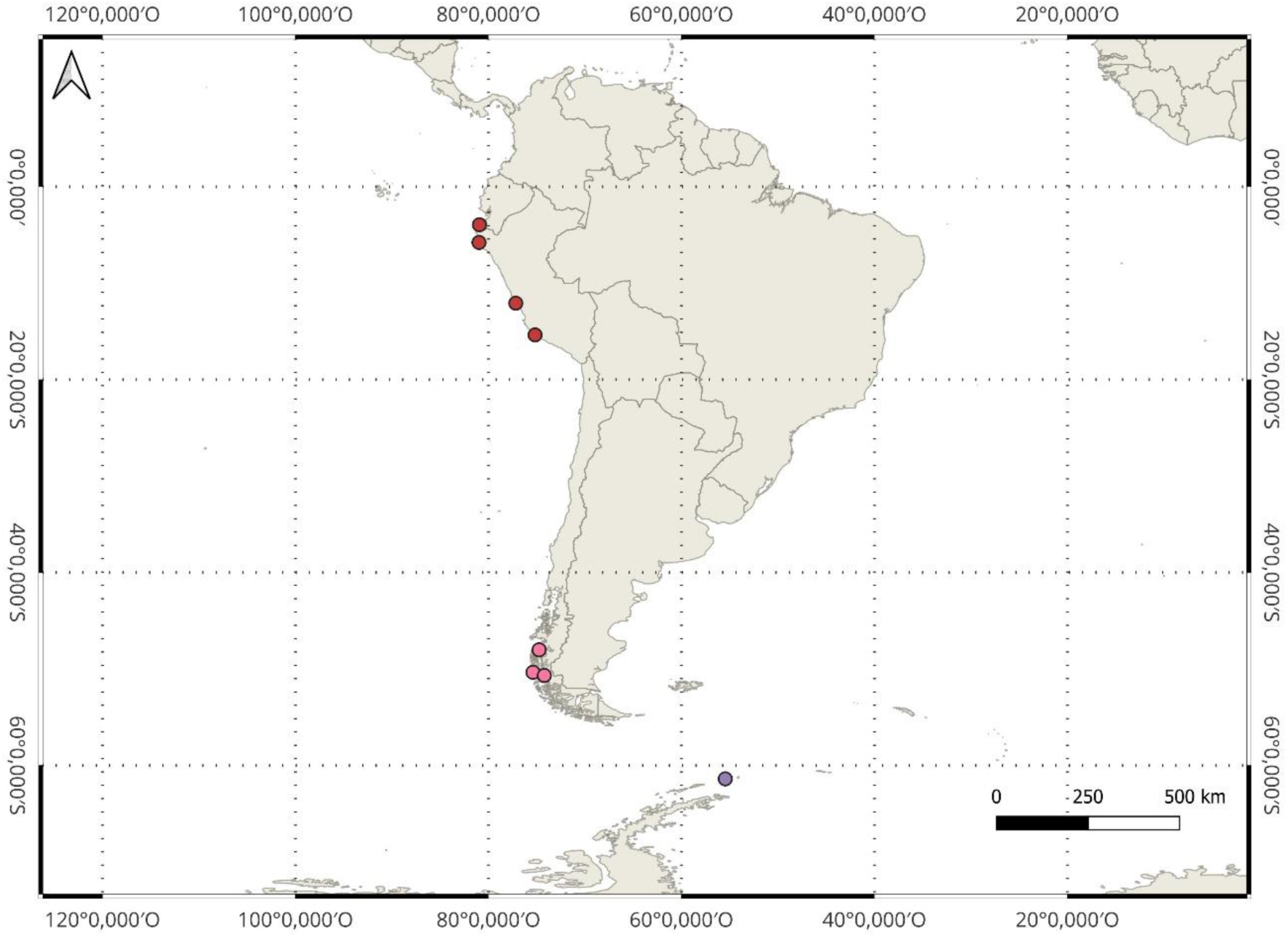
Distribution map from the South America and Antarctic Peninsula of the *Doto* species sequenced in this study. Circles represent sampling stations colour-coded by region: Peru (red), Chile (pink), South Shetland Islands, Antarctica (purple). Map generated using QGIS v.3.36.0

### DNA extraction, amplification, and sequencing

Genomic DNA was extracted from a few dorsolateral appendages using the Speedtools Tissue DNA Extraction kit (Biotools), according to the manufacturer’s protocol. Sequences of three standard molecular markers were amplified, i.e., the mitochondrial cytochrome c oxidase subunit I (COI; ca. 658 bp) using the LCO1490 and HCO2198 primer pair (Folmer et al., 1994) and 16S rRNA (16S; ca. 485 bp) using the primers 16S ar-L and 16S br-H (Palumbi et al., 1991); and the nuclear gene histone H3 (H3; ca. 330 bp) with H3AD5’3’ and H3BD5’3’ (Colgan et al., 1998). Polymerase chain reactions (PCR) were carried out in 20 μL volume reactions with 9 μL of Sigma dH2O, 8 μL of 5x MyTaqTM Red DNA Polymerase, 0.5 μL of each primer, and 2 μL of genomic DNA of each sample. PCR programs for COI involve an initial denaturation step (95 °C for 5 min), followed by 35 cycles of denaturation (95 °C for 30 s), annealing (45–55 °C for 35 s), and extension (72 °C for 45 s), with a final extension step at 72 °C for 5 min. Regarding 16S, a hot start step of 15 min at 95°C was followed by 40 cycles of 30 s at 94 °C, 90 s at 46–52 °C, 90 s at 72 °C, and a final extension for 10 min at 72 °C. The conditions for the H3 marker differed in that the annealing temperature was set to 55 °C. Successful amplifications were sequenced by Macrogen, Inc. (Madrid, Spain) after ExoSAP-IT™ Express PCR Product Cleanup Reagent purification. Sequences were uploaded to GenBank, and accession numbers are provided in Table S1.

### Phylogenetic analyses

Potential contaminations were checked for each sequence against the GenBank nucleotide database using the BLAST algorithm (Altschul et al., 1997). Sequences were visualized, edited, and assembled in Geneious Prime® 2023.0.4 (Kearse et al., 2012). MAFFT v 7.450 was used for multiple alignments (Kato & Standley, 2013), using G-INS-i (COI and H3) and L-INS-i (16S) algorithms.

Phylogenetic analyses, both for single-gene alignments and concatenated genes, involved a maximum likelihood (ML) approach implemented in IQ-TREE v. 2.1.2. (Nguyen et al., 2015). The best substitution model was automatically selected using ModelFinder (Kalyaanamoorthy et al., 2017) for each partition and accounting for codon positions. The TESTMERGE option was used to reduce overparameterization and increase model fit.

Bootstrap support values (bs) were estimated via the ultrafast bootstrap algorithm with 1,500 replicates (Hoang et al., 2018). A Bayesian inference (BI) analysis was performed with the concatenated alignment using MrBayes v 3.2.7 (Ronquist et al., 2012), applying the GTR+G+I evolution model per partition. Four parallel runs of four coupled Markov Chain Monte Carlo (MCMC) chains were run for 20 million generations, sampling every 1,000. The first 25% of the trees were discarded as burn-in for each MCMC run before convergence.

Convergence was achieved when the average standard deviation of split frequencies reached <0.01% for all parameters. Topological robustness was assessed using posterior probabilities (pp). Trees were visualized in FigTree v. 1.4.4 (Rambaut, 2014) and edited in Adobe Illustrator.

Species delimitation tests (SDT) were conducted on the COI alignment only. The Assemble Species by Automatic Partitioning (ASAP; Puillandre et al., 2021) analysis was run using the web interface at https://bioinfo.mnhn.fr/abi/public/asap/, applying a Kimura (K80) model of sequence evolution with TV/TS = 2.0. Moreover, the Poisson Tree Processes (PTP, Zhang et al., 2013) was run using the web interface at mPTP Webservice (https://mptp.h-its.org/#/tree).

### Anatomical analyses

Detailed descriptions of the external morphology and egg masses of 15 species are provided based on underwater photographs of living specimens, laboratory images, and bibliographic descriptions. To aid in differentiating among the Atlanto-Mediterranean species, comparative schematic illustrations of 13 species were depicted.

Animal length (L) was measured with the aid of a micrometric ruler. Dissections of the newly described species were carried out: (1) the reproductive system was carefully removed to be observed and accurately described; (2) the buccal bulb was extracted, immersed in a 10% KOH solution for 1 hour (to dissolve organic tissues), and then rinsed with distilled water. This was followed by an ultrasonic bath (to remove the remaining organic tissue) with three dips for 5 seconds with distilled water. The radula was mounted on metal slides with carbon adhesive tabs and coated with carbon for scanning electron microscopy (SEM) with a Field Emission Scanning Electron Microscope JSM-7100F at the UB Scientific and Technological Centres (CCiT-UB).

## Results

### Phylogenetic assessment

The total dataset included 169 samples of *Doto* and two outgroup species of the closest genus *Kabeiro* (Dotidae), according to Moles et al. (2016). Out of these 171 specimens, 59 were sequenced for this study, while the rest were downloaded from GenBank (see Table S1). The final concatenated alignment contained 1423 bp, with 658 bp for COI (Fig. S1), 437 characters for 16S (Fig. S2), and 328 bp for H3 (Fig. S3). The best-fit model of sequence evolution was TVM+F+I+G4 for all partitions and codon positions, except for H3, which was TPM3+F+I+G4. Both ML (Fig. 3) and BI (Fig. S4) yielded similar topologies for the concatenated alignment.

**Fig. 3.**
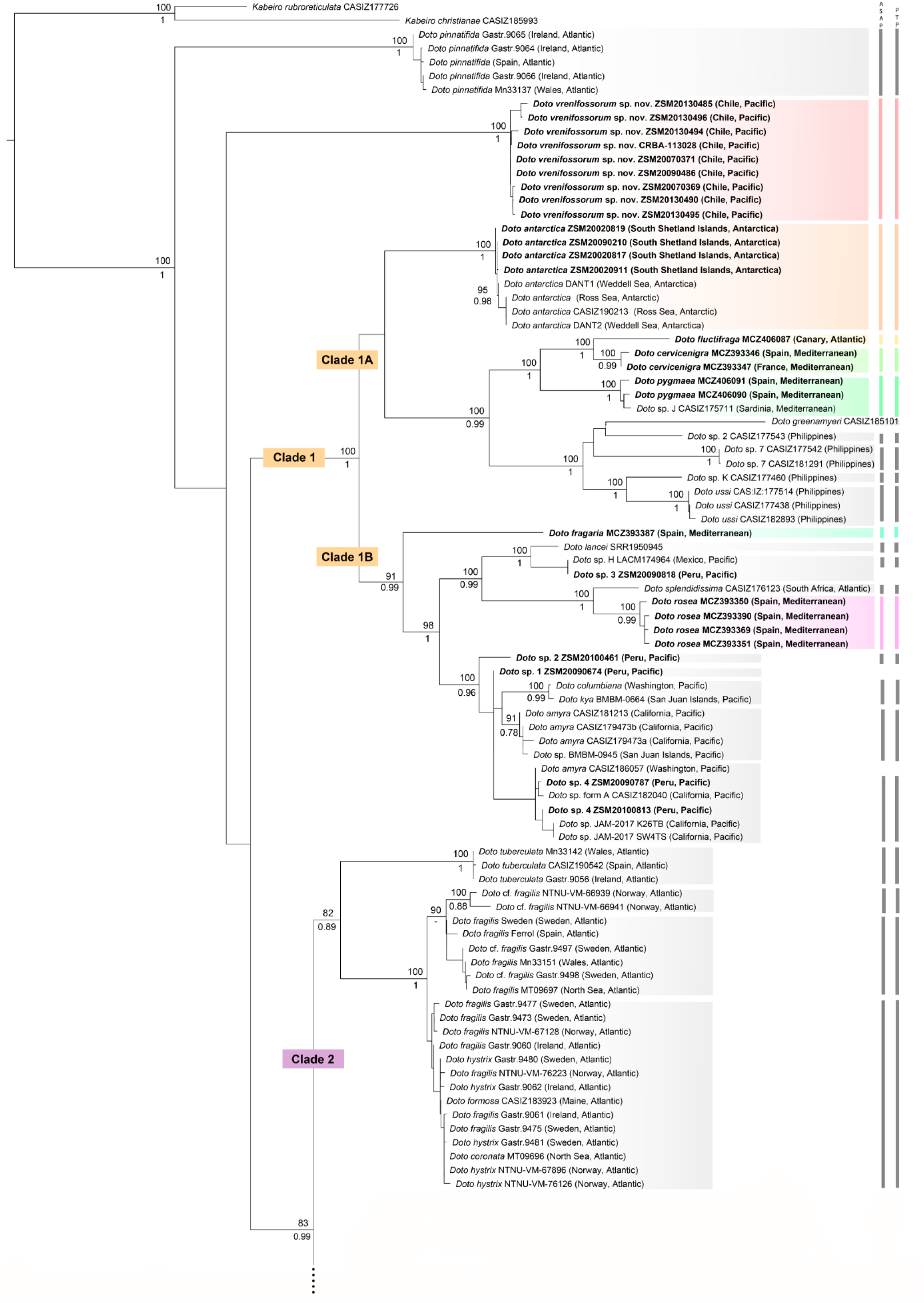
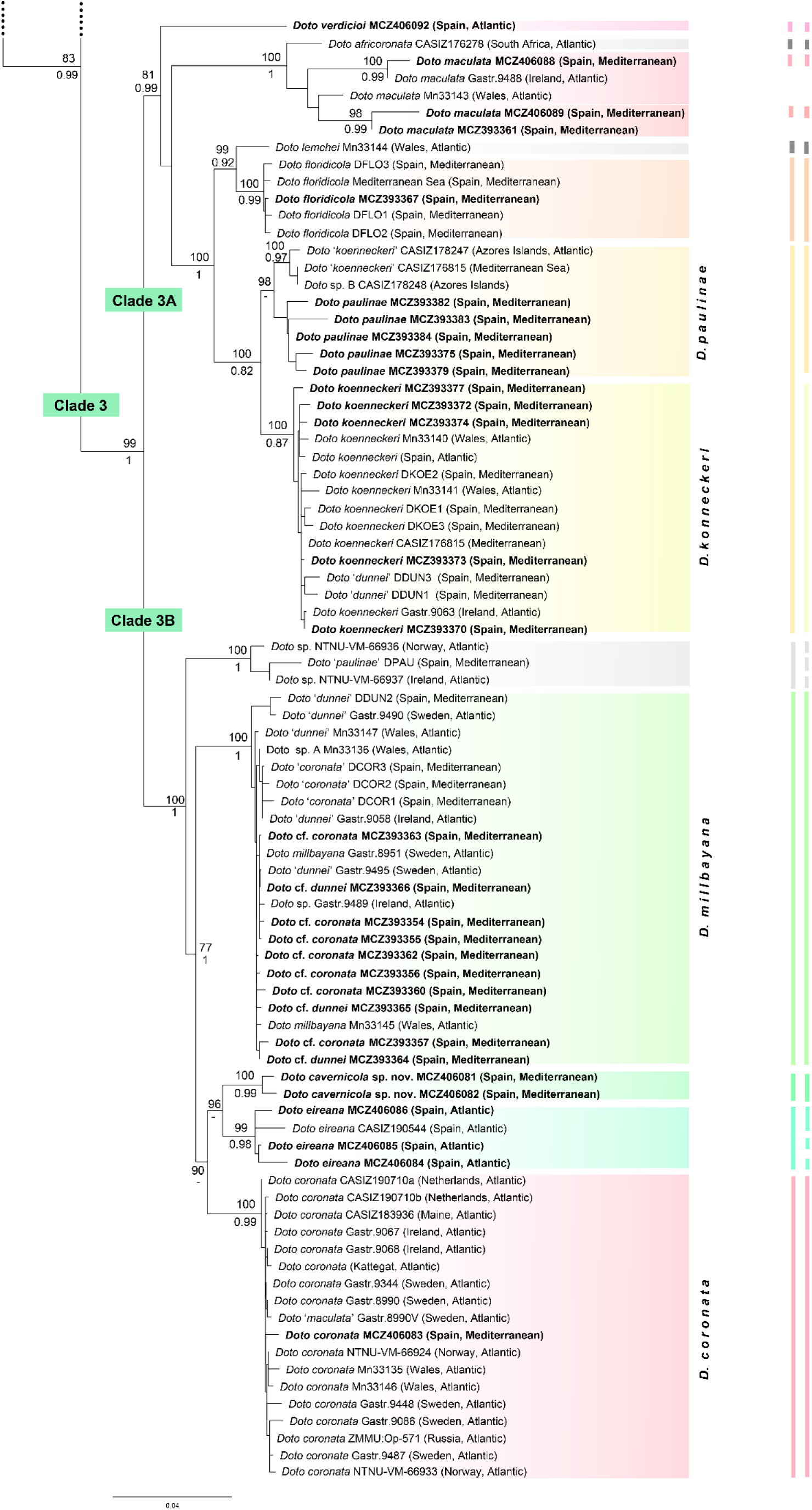
Phylogenetic tree of *Doto* species constructed with maximum likelihood based on the concatenated mitochondrial (COI and 16S) and nuclear (H3) markers. Support values shown on branch nodes are based on bootstrap support values (ML) above nodes and posterior probabilities (BI) below. Specimens in bold were newly sequenced for this study. The results of ASAP and PTP species delimitation analyses based on COI are represented by the vertical bars to the right side and by the coloured boxes. The scale bar represents substitutions per site. Only values above 0.78 for BI posterior probabilities and 75 for ML bootstraps are depicted.

The first clade (Fig. 3) consists of *D. pinnatifida* (bs = 100, pp = 1.00). A second clade includes nine new specimens from South America corresponding to an undescribed species, here described as *D. vrenifossorum* sp. nov., with maximum support in both analyses. Nevertheless, the relationships between *D. pinnatifida*, *D. vrenifossorum* sp. nov., and other large clades are not supported by both analyses.

All other *Doto* species are encompassed into three clades labelled in Fig. 3. Clade 1 (bs = 100, pp = 1.00) can be subdivided into two groups, Clade 1A, which lacks support, comprising *D. antarctica* Eliot, 1907 from the Weddell and the Ross seas, along with four new specimens from the South Shetland Islands; three species (bs = 100, pp = 1.00) sequenced for the first time from the Canary Islands (*D. fluctifraga* Ortea & Perez, 1982) and the Mediterranean Sea (*D. cervicenigra* Ortea & Bouchet, 1989 and *D. pygmaea* Bergh, 1871); as well as specimens from the Papua New Guinea (*D. greenamyeri*) and the Philippines (*D. ussi*, and other unidentified species) (bs = 100, pp = 1.00). Clade 1B (bs = 91, pp = 0.99) comprises *D. fragaria* Ortea & Bouchet, 1989 from the Mediterranean Sea, sequenced here for the first time; a second group (bs = 100, pp = 0.99) contained *D. lancei* Ev. Marcus & Er. Marcus, 1967, two unidentified species from Mexico and Peru, *D. splendidissima* Pola & Gosliner, 2015 from South Africa, and *D. rosea* Trinchese, 1881 from the Mediterranean; and several species (bs = 100, pp = 0.96) from the Pacific of North America (*D. columbiana* O’Donoghue, 1921, *D. kya* Er. Marcus, 1961, *D. amyra* Er. Marcus, 1961, and other unidentified species), along with our four newly sequenced specimens from Peru.

Clade 2, which has low support, encompasses the species *D. tuberculata*, *D. fragilis* (Forbes, 1838), *D. formosa* A. E. Verrill, 1875, and *D. hystrix* from the NE Atlantic (bs = 82, pp = 0.89). All of them were obtained from GenBank and their relationships did not differ from previous studies (Martinsson et al., 2021).

Clade 3 includes species from the NE Atlantic and the Mediterranean Sea (bs = 99, pp = 1.00). Clade 3A (bs = 81, pp = 0.99) contains *D. verdicioi* from the Cantabric Sea (NE Atlantic Ocean), sequenced here for the first time, as the first offshoot. Another clade is composed of *D. africoronata* Shipman & Gosliner, 2015 from South Africa and *D. maculata* from the NE Atlantic and the Mediterranean Sea (bs = 100, pp = 1.00). These are the first confirmed records of *D. maculata* in the western Mediterranean, although the monophyly of *D. maculata* is not supported. The last clade within 3A (bs = 100, pp = 1.00) is composed of *D. lemchei* as sister group to *D. floridicola* (bs = 99, pp = 0.92), and these being sister to the clade composed of *D. koenneckeri* and *D. paulinae* (bs = 100, pp = 0.82). Within the latter clade, two specimens identified as *D. koenneckeri* from the Mediterranean and Azores, along with unidentified species from the Azores (bs = 100, pp = 0.97), are grouped with *D. paulinae* only in the ML analysis (bs = 98), while in the BI they appeared scrambled all over albeit without statistical support. In the same clade, two specimens identified as *D. dunnei* from the Mediterranean were found to group with *D. koenneckeri* (bs = 100, pp = 0.87). These may be misidentifications according to our results.

Clade 3B (bs = 100, pp = 1) is composed of a first splitting branch with three unidentified specimens from the NE Atlantic and W Mediterranean. A second group includes *D. millbayana* and *D. dunnei* from the Mediterranean Sea and Atlantic Ocean and the supposed *D. coronata* specimens from the Mediterranean (bs = 100, pp = 1.00) (see discussion below). Moreover, we obtained a clade of the *D. coronata* species complex (bs = 90). The latter includes *D. coronata* from the type locality in the Netherlands and our only newly sequenced specimen from the Mediterranean (bs = 100, pp = 0.99). Additionally, two similar-looking species are also included in the *D. coronata* species complex (bs = 96). These are the Mediterranean specimens found in caves here newly described as *Doto cavernicola* sp. nov. (bs = 100, pp = 0.99) and *D. eireana* composed of three newly sequenced specimens from Galicia, NW Spain (bs = 99, pp = 0.98).

The ASAP and PTP analyses support most of the abovementioned results (see Fig. 3), although some discrepancies are present. In both SDTs, *D. columbiana* and *D. kya* were clustered together, *D. fragilis* was divided into three groups, and *D. amyra* into two. On the other hand, in the ASAP analysis, *D. paulinae* and *D. koenneckeri* were clustered together, while the PTP analysis delimited the specimens of *D. eireana* into three different clusters, as well as the unidentified specimens from the NE Atlantic and the W Mediterranean (*Doto* sp. / *paulinae*).

### Systematic descriptions

Subclass Heterobranchia Burmeister, 1837

Order Nudibranchia Cuvier, 1817

Family Dotidae Gray, 1853

Genus Doto Oken, 1815

***Doto cavernicola* sp. nov. Vázquez-Alcaide, Salvador & Moles, 2025**

(Fig. 4, Fig. 5b)

**Fig. 4.**
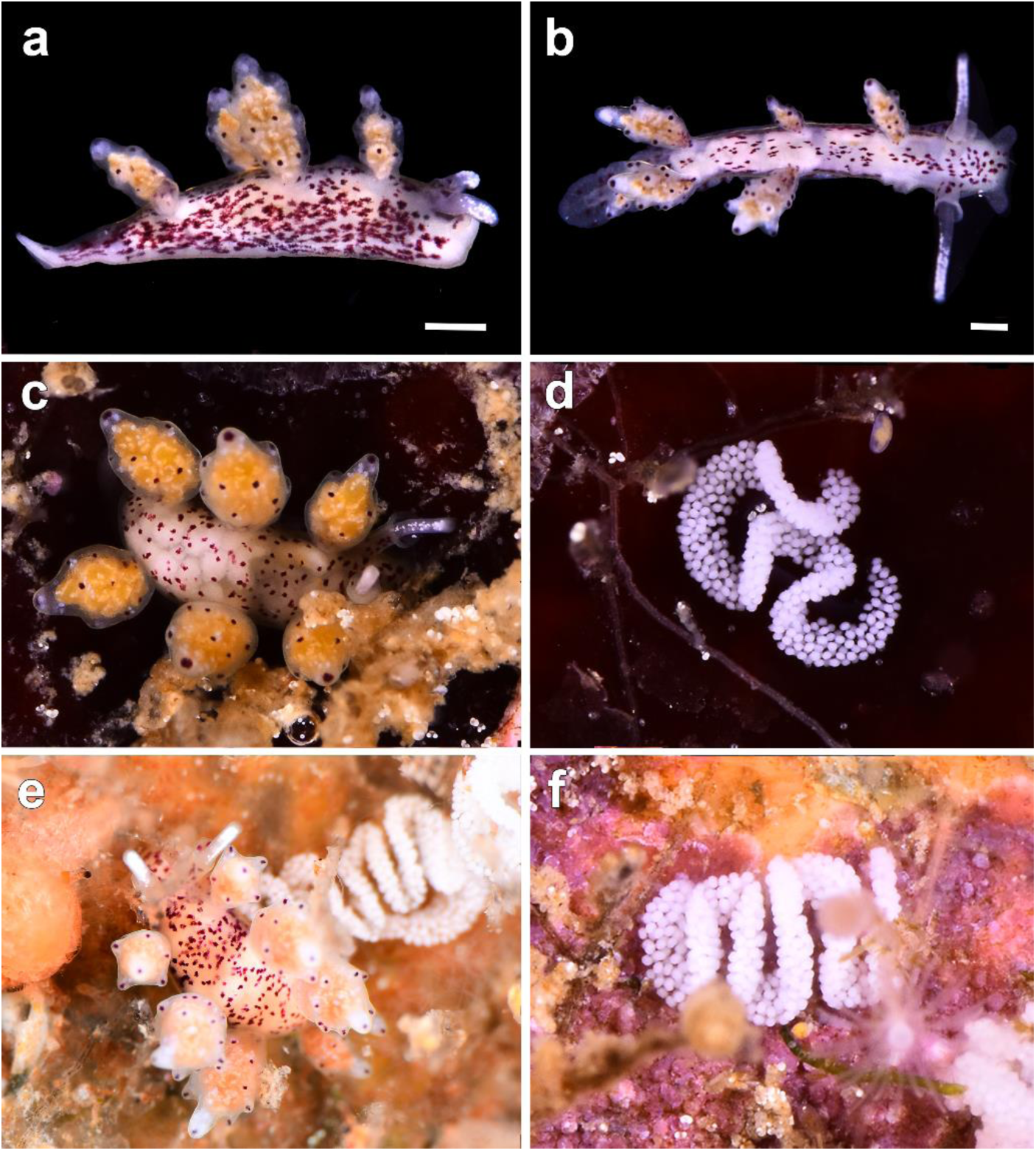
Photographs of living specimens of the new species *Doto cavernicola* sp. nov. **a** holotype MCZ:Mala:406082 **b** paratype MCZ:Mala:406081 **c** specimen found living in a cave entrance **d** the egg mass on a rocky surface **e** specimen with their egg mass fixed on the hydroid *Diphasia* cf. *delagei* **f** egg mass fixed on a rock. Scale bar: 0.2 mm. Photos by Xavier Salvador

**Fig. 5.**
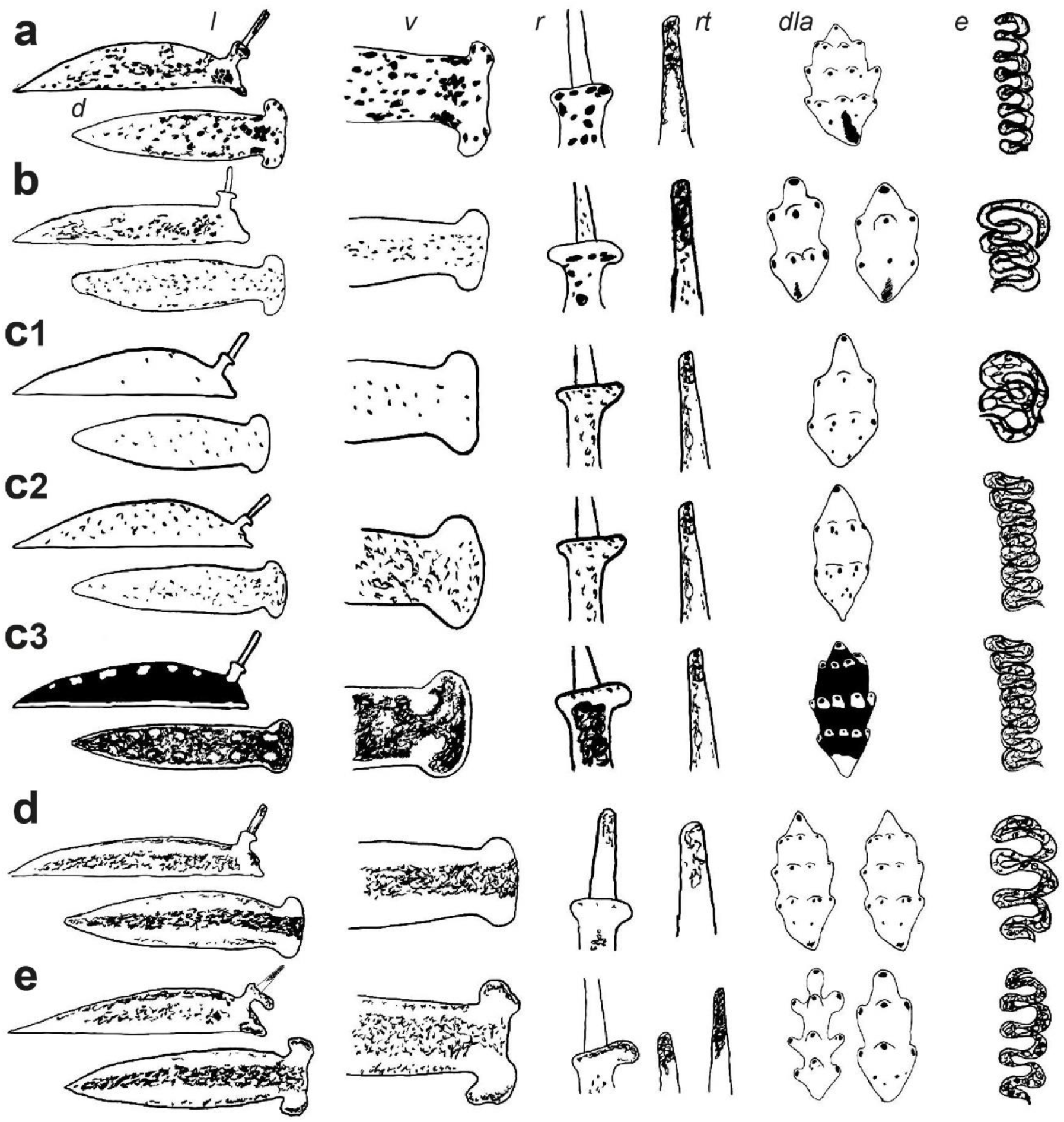
Illustrations of the external morphology of the Mediterranean *Doto* species included in this study. The different species indicated in lowercase bold letters **a** *D. coronata* **b** *D. cavernicola* sp. nov. **c1** *D. millbayana* **c2** *D. millbayana* morphotype traditionally attributed to *D. coronata* **c3** *D. millbayana* morphotype traditionally attributed to *D. dunnei* **d** *D. eireana* **e** *D. maculata*. Parts of the external morphology indicated in lowercase italics letters. d dorsal view; l lateral view; v oral veil; r rinophores; rt rinophores tip; dla dorsolateral appendages; e egg mass

**ZooBank registration link.** https://zoobank.org/NomenclaturalActs/c51175ba-a4c7-4474-ae37-155c636e1168

**Type material examined**. ***Holotype.*** SPAIN · L = 2.5 mm (sequenced); Catalonia, Girona, Sant Feliu de Guíxols, Cala Maset caves; 41°47’10.5“N, 03°02’44.6”E, 1 m depth; 06 May 2020; X. Salvador leg.; MCZ:Mala:406082. ***Paratypes*.** SPAIN · 4 spcs, L = 1.5–2 mm (one sequenced and two dissected); same collecting place and date as for holotype; X. Salvador leg.; MCZ:Mala:406081.

**Etymology**. The species is named after the habitat where it was found, the entrance of submarine caves.

**Diagnosis**. Body with irregular red dots in dorsum, more abundant in laterals and without dots around dorsolateral appendages. Rhinophores white pigmented in apical half. Rhinophoral sheath internally pigmented with some red dots at base and apex, visible through background transparency. Dorsolateral appendages in 3–4 pairs; tubercles in 2–3 crowns, extended, with one red apical dot; base pigmented with irregular dots. Oral veil unpigmented.

**Description.** (Fig. 4, Fig. 5b) Body elongated, narrow, background white, with irregular red dots scattered all over, except for dorsum around dorsolateral appendages. Rhinophores white pigmented in apical half; rhinophoral sheath short, base and apex pigmented with red dots internal and posteriorly, seen internally by transparency in margin. Dorsolateral appendages in 3–4 pairs; tubercles in 2–3 crowns, extended, with one red, apical dot; base pigmented with irregular dots. Oral veil unpigmented.

**Radula.** The radula could not be recovered from the dissected specimens.

**Reproductive system.** (Fig. 6) Ampulla sausage-shaped; from the distal part a thin gonoduct extends into prostate and oviduct. ♂ Prostate long, folded, attached to a thin distal vas deferens, connected to penis. Penis short, ovoid; penial sheath saccular. Genital openings situated in anterior-right position. ♀ Oviduct branched, distally towards seminal receptacle and nidamental glands. Seminal receptacle small, rounded, saccular, attached to vaginal canal. Nidamental glands large, sharing a wide atrium with vagina. Vagina short, flattened, reaching the exterior through a wide opening.

**Fig. 6.**
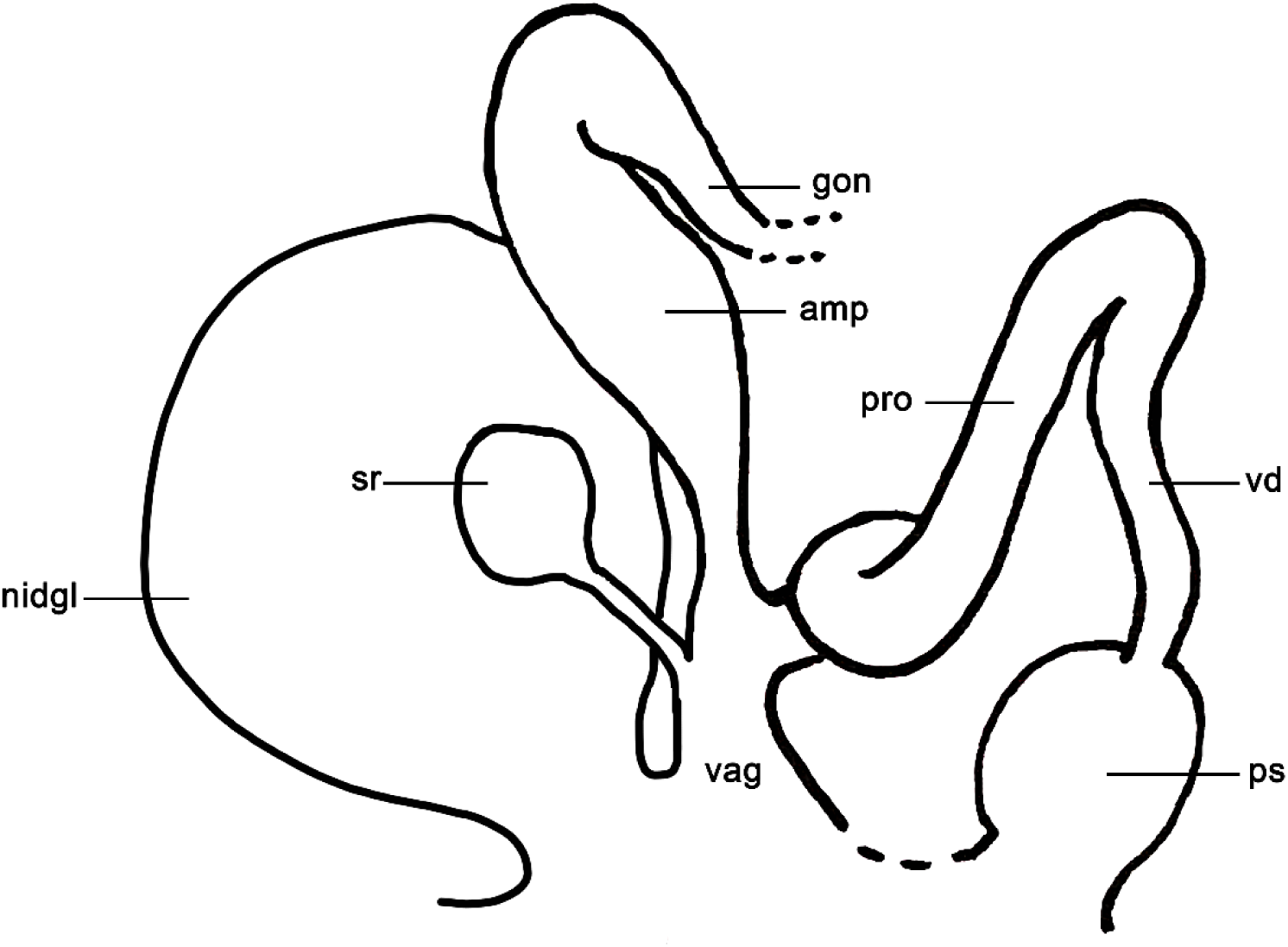
Schematic drawing of the reproductive system of *Doto cavernicola* sp. nov. Abbreviations: amp, ampulla; gon, gonoduct; nidgl, nidamental glands; ps, penial sheath; pro, prostate; sr, seminal receptacle; vag, vagina; vd, vas deferens

**Egg mass.** (Fig. 4d, 4f, Fig. 5b) White, displayed in a meandering cord shape. The egg masses are usually fixed on rocks or over hydrozoans, with up to 3–5 folds.

**Distribution.** Only known from the Catalan coast (Mediterranean Sea).

**Ecology.** The specimens were found on rocky substrates in the entrance of shallow waters caves in different diving areas at 1 m depth, always living at the base of a stoloniferous hydrozoans *Diphasia* Agassiz, 1862, of the species *Diphasia* cf. *delagei*. They are more active during the night, hiding in the base of the hydrozoans during the day.

**Life cycle.** The first specimens were found from December until June. The first egg masses were found in February, with a second period of egg laying in May–June. Subsequently, with the raise of temperature the hydrozoans die, and the nudibranchs end their life cycle.

**Remarks.** Morphologically, *D. cavernicola* sp. nov. is much alike *D. coronata*. They appear in the same clade along with *D. eireana*. Diagnostic features of *D cavernicola* sp. nov. are its diet, as it feeds on *Diphasia* cf*. delagei*, the absence of dots around the dorsolateral appendages and the unpigmented oral veil, while *D. coronata* from the Mediterranean has larger dots on the body and dorsolateral appendages, the base of the rhinophores is more pigmented and the oral veil is pigmented. Instead, *D. eireana* usually lacks apical dots on the apical tubercle of dorsolateral appendages. Moreover, the habitat where the specimens were found and the shape of the egg mass, so far, are characteristic of the new species. The reproductive system of *D. cavernicola* sp. nov. also differs from that of *D. coronata*. In *D. cavernicola* sp. nov. the ampulla is sausage-shaped, the vagina is short and flattened, and the penis and penial sheath are short, while in *D. coronata* the ampulla is shaped like ‘a human stomach’, the vagina is elongated and narrow, and the penis and penial sheath are more elongated (Schmekel & Portmann, 1982; Shipman & Gosliner, 2015). In the eastern North Atlantic, Morrow et al. (1992) described *D. hydrallmaniae* and *D. sarsiae* under the umbrella of the *D. coronata* complex. Molecular data do not validate both species and exhibit external morphological similarities to *D. cavernicola* sp. nov.

Unlike *D. cavernicola* sp. nov, *D. hydrallmaniae* has larger dark red pigment spots on the tubercles, which may fuse to form irregular marks between them. It also presents more elongated tubercles with a larger number of crowns (Picton & Morrow, 2023). Regarding *D. sarsiae*, the species also has larger dark red spots on the tubercles and body, the digestive gland is pinkish red, and some specimens lack apical dots on the apical tubercle (Picton & Morrow, 2023). Additionally, their distribution and diet are diverse, *D. hydrallmaniae* feeds on *Hydrallmania falcata* (Linnaeus, 1758) and *D. sarsiae* on *Coryne eximia* Allman, 1859, and both are distributed in the North-East Atlantic (Morrow et al., 1992; Picton & Morrow, 2023).

***Doto cervicenigra* Ortea & Bouchet, 1989**

(Fig. 7, Fig. 8g)

**Fig. 7.**
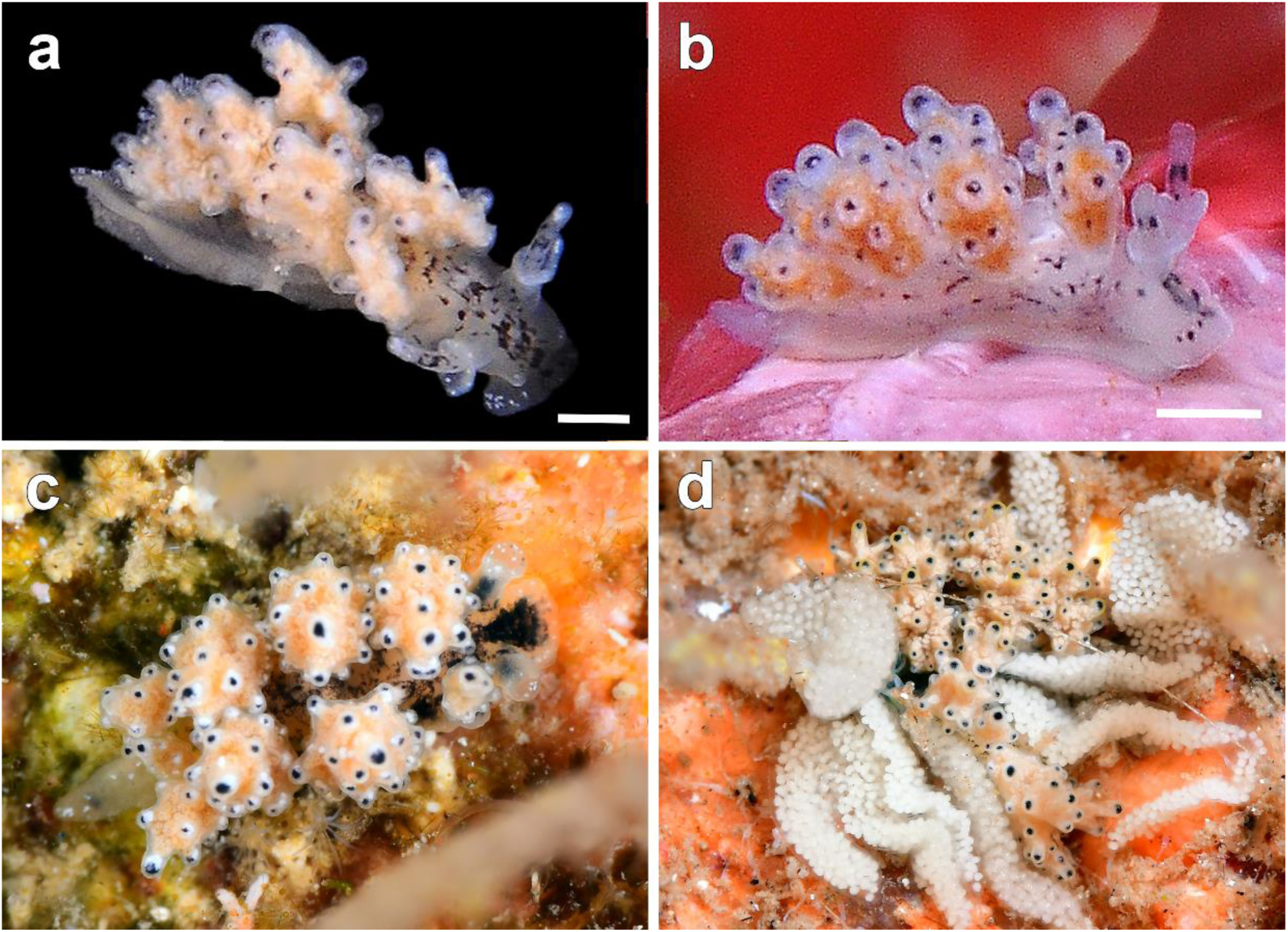
Photographs of live *Doto cervicenigra* specimens. Sequenced specimens **a** MCZ:Mala:393347 and **b** MCZ:Mala:393346 **c–d** *D. cervicenigra* and its egg mass. Scale bar: 1 mm. Photos by Xavi Salvador

**Fig. 8.**
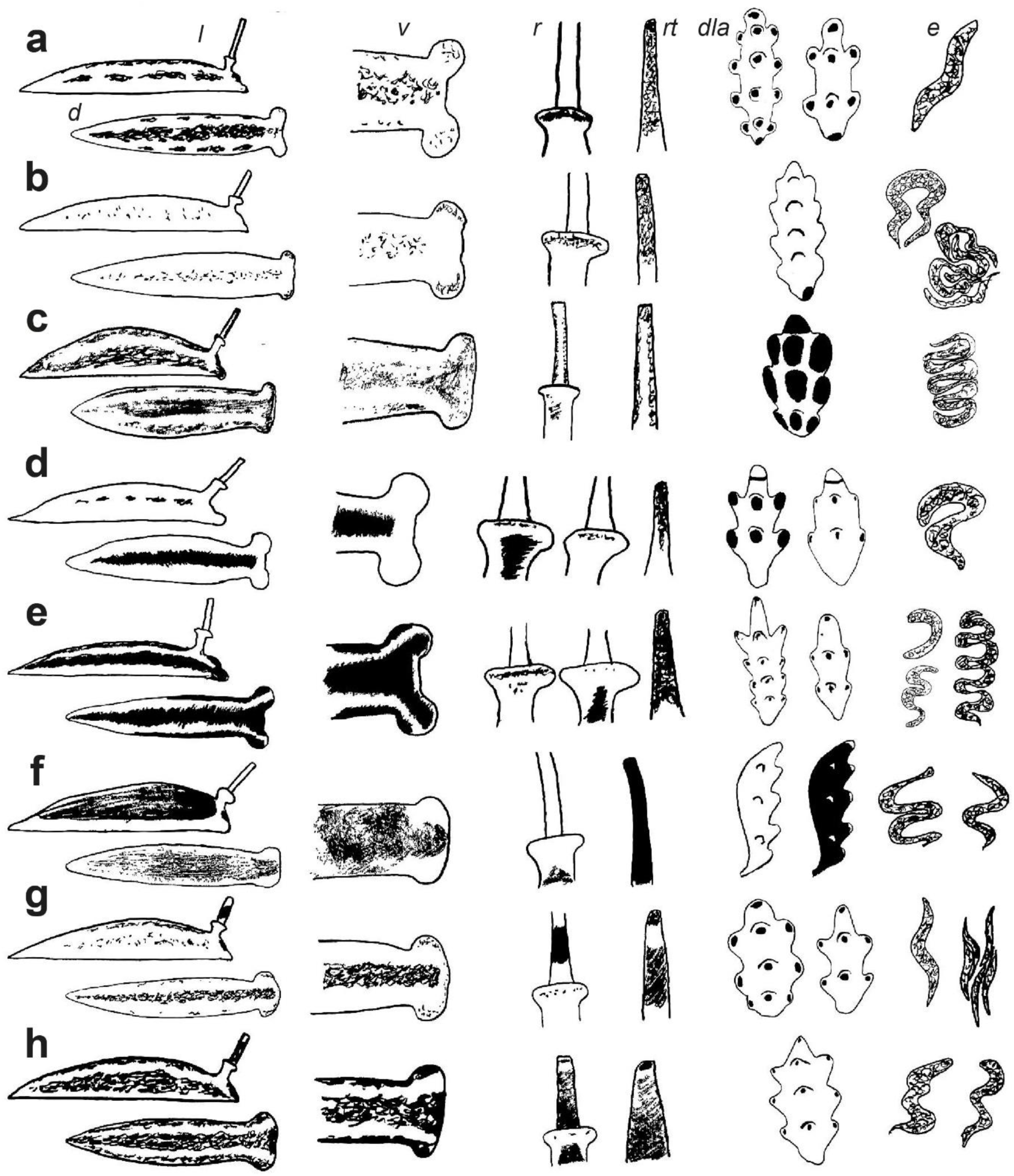
Illustrations of the external morphology of the Mediterranean and North Atlantic *Doto* species included in this study. The different species indicated in lowercase bold letters **a** *D. fragaria* **b** *D. rosea* **c** *D. floridicola* **d** *D. paulinae* **e** *D. koenneckeri* **f** *D. pygmaea* **g** *D. cervicenigra* **h** *D. fluctifraga*. Parts of the external morphology indicated in lowercase italics letters. d dorsal view; l lateral view; v oral veil; r rinophores; rt rinophores tip; dla dorsolateral appendages; e egg mass

*Doto cervicenigra* Ortea & Bouchet, 1989: 265–268; Ballesteros, Madrenas & Pontes, 2016: 12; Lombardo, 2021: 916–917; Salvador, Fernández-Vilert & Moles, 2022: 279–280.

**Material examined.** SPAIN · 1 spc., L = 5 mm (sequenced); Catalonia, Girona, Sant Feliu de Guíxols, Cala Maset caves; 41°47’10.5“N, 03°02’44.6”E, 1 m Depth; 13 Jan. 2018; X. Salvador leg.; MCZ:Mala:393346 · FRANCE · 1 spc., L = 7 mm (sequenced); Sete, étang de Thau; 43°25’48“N, 03°42’14”E, 0.2 m depth; 20 May 2018; X. Salvador leg.; MCZ:Mala:393347.

**External morphology.** (Fig. 7, Fig. 8g) Body elongate, translucent, white background; black irregular markings on head extending backwards between rhinophores; black irregular markings scattered on surface, densely interconnected. Rhinophores with black, central spots, white apically; rhinophoral sheath short, thin, symmetrical, with barely flared anterior edge, and mottled with opaque white dots. Dorsolateral appendages arranged in 5–6 pairs; tubercles displayed in 2–3 crowns, almost spherical, with black dots on tip.

**Egg mass.** (Fig. 7d, Fig. 8g) Saccular, short, elongated and with straight shape; translucent in color, with white eggs.

**Distribution**. France: Corsica (Ortea & Bouchet, 1989); Italy (Ballesteros et al., 2012–2024): Sardinia (Trainito & Donneddu, 2015), Sicily (Lombardo & Marletta, 2020), and western Adriatic Sea (Riccardi et al., 2022); Spain: Catalonia (Ballesteros et al., 2016), Mallorca (GROC, 2024); France (Salvador, Fernández-Vilert & Moles, 2022).

**Ecology.** Both specimens were found at shallow depths, crawling on hydrozoan species of probably the genus *Obelia* Péron & Lesueur, 1810, *Sertularella mediterranea* Hartlaub, 1901, and *Campanularia* Lamarck, 1816, where they were feeding and laying the egg masses at the bottom of the colony.

**Life cycle.** Adults are found between October and May; mating behavior and egg masses were recorded between January and May (based on 124 records of the GROC database, including 35 sampling dates and 11 different localities).

**Remarks.** *D. cervicenigra* is the only species in the Mediterranean known to have black pigmentation on the rhinophores. These characteristics make it easily distinguished from other sympatric *Doto* species. In the North Atlantic Ocean, *D. eo* Ortea & Moro, 2014 and *D. fluctifraga* also present black pigmentation on the rhinophores. *Doto eo* differs from *D. cervicenigra* by the salmon colour of the dorsolateral appendages and the conical and less prominent shape of their tubercles (Ortea & Moro, 2014). As for *D. fluctifraga*, no external morphological differences from *D. cervicenigra* are evident although confirmed as two different species according to the SDT. Thus, we rely on the differences in their distribution and diet. The Atlantic *D. fluctifraga* feeds on *Pennaria disticha* Goldfuss, 1820, while the Mediterranean *D. cervicenigra* feeds on *Obelia*, *Campanularia*, and *Sertularella mediterranea*.

***Doto coronata* (Gmelin, 1791)**

(Fig. 9, Fig. 5a)

**Fig. 9.**
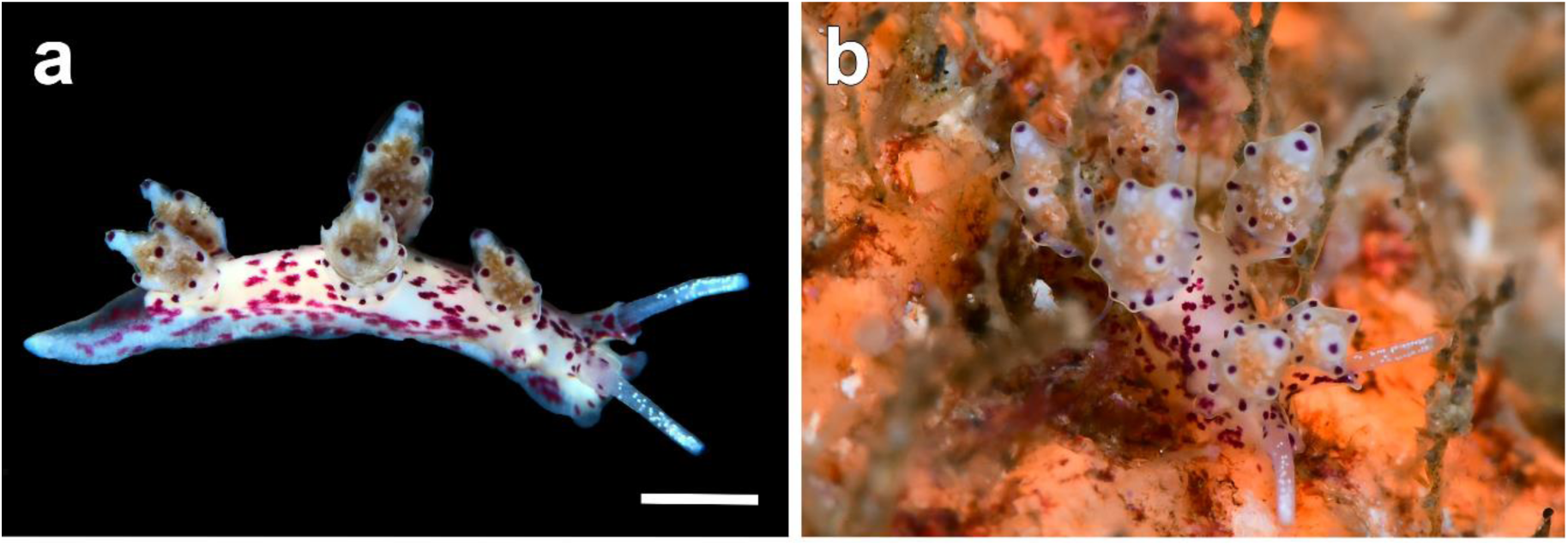
Photographs of live *Doto coronata* specimen **a** *D. coronata* (MCZ:Mala:406083) **b** same specimen between colonies of the hydroid. Scale bar: 1 mm. Photos by Xavi Salvador

*Doris coronata* Gmelin, 1791: 3021–3910.

*Doto coronata*: Lemche, 1976: 694–695; Vogel, 1978: 74–76; Schmekel & Portmann, 1982: 158–168; Thompson, Cattaneo & Wong, 1990: 394–396; Ortea et al., 2008: 1333; Shipman & Gosliner, 2015: 24–28;

Picton & Morrow, 2023: 170–171.

*Doto costae* Trinchese, 1881:91.

*Doto forbesii* Deshayes, 1853: original description not documented.

*Doto splendida* Trinchese, 1881:93.

**Material examined.** SPAIN · 2 spcs, L = 5 mm (one sequenced); Catalonia, Girona, Cala Maset caves; 41°47’10.5“N, 03°02’44.6”E, 1 m depth; 17 Febr. 2022; X. Salvador leg.; MCZ:Mala:40608.

**External morphology.** (Fig. 9, Fig. 5a) Body elongated, narrow, translucent, cream to white background, with red to maroon irregular dots scattered all over, except for dorsum around dorsolateral appendages. Oral veil pigmented. Rhinophores white pigmented in apical half; rhinophoral sheath cylindrical, symmetrically, with red punctuation at base and margin. Dorsolateral appendages arranged in 3–5 pairs, wide, base pigmented of reddish-maroon irregular dots; tubercles displayed in 2–3 crowns, apical part pointed, with single red dot.

**Egg mass.** (Fig. 5a) No egg masses were found associated with the collected specimen. The egg mass of Atlantic specimens is described as a cream-coloured, double-looped ribbon deposited along the stem of the hydroid (Lemche, 1976; Shipman & Gosliner, 2015).

**Distribution.** Amphi-Atlantic, in the western Atlantic from New England, New Jersey, and Long Island (Verrill & Smith, 1874; Shipman & Gosliner, 2015); in the eastern North Atlantic from Iceland and Scandinavia to the Iberian Peninsula (Thompson et al., 1990); in the Barents Sea (Martynov et al., 2006); reaching the Canary Islands (Ortea et al., 2008); Catalan coast (in this study).

In the Mediterranean Sea some citations could be misidentified with *D. millbayana*, further records from Catalonia (Thompson et al., 1990); Bay of Naples (Schmekel & Kress, 1977); Aegean and Adriatic Seas (Odhner, 1914); Southeastern France (Vicente, 1967); Turkish coasts (Öztürk et al., 2014), Malta (Sammut & Perrone, 1998), and Italy (Ballesteros et al., 2012–2024), need reassessment.

**Ecology.** The Mediterranean specimens were found at a cave entrance, in vertical rocky walls covered by hydrozoans of the genus *Sertularia* Linnaeus, 1758 [probably *Sertularia distans* (Lamouroux, 1816)].

Specimens were found more active at night.

**Life cycle.** Given the present reclassification of the Mediterranean species of *D. coronata* and *D. dunnei* (see Phylogenetic results), ascertaining the life cycle of this species requires additional sampling. It is observed in the winter, particularly in February, along the Catalan coast.

**Remarks.** In the original description of *D. coronata* it is mentioned the presence of a single spot at each tubercle of the dorsolateral appendages, the pigmented base, and the whole body with plenty of markings (Gmelin, 1791), from red to black (Picton et al., 2010). Many specimens from the Mediterranean that fit that description are here described as *D. millbayana*, which encompasses a broad chromatic variation (see description below). These are particularly found in *Posidonia* meadows (this study) and are usually wrongly attributed to *D.* ‘*coronata*’. The few specimens of *D. coronata* confirmed with molecular data in the Mediterranean present a single dot in each tubercle and lack markings between them, while *D. millbayana* has more than one spot per tubercle and many markings between them, especially in the lower part. Ecologically, the Mediterranean specimens correctly identified as *D. coronata* lived in the entrance of caves and walls with high hydrozoan coverage (*Sertularia*). Also, specimens of *D. coronata* were found to be more active at night, while *D. millbayana* is especially active during daylight.

In the Canary Islands (eastern Atlantic), *D. canaricoronata* Moro & Ortea, 2015 can also be confused with *D. coronata*. This species can be distinguished from *D. coronata* by its white dorsolateral appendages, rings of red tubercles arranged in a linear fashion, and the absence of reddish pigmentation at the base of the dorsolateral appendages (Moro & Ortea, 2015). Given the morphological diversity within of *D. coronata* species complex, conducting a comparative analysis of *D. coronata* and *D. canaricoronata*, including molecular data, would serve to reinforce the validity of this species.

**Doto eireana Lemche, 1976**

(Fig. 10, Fig. 5d)

**Fig. 10.**
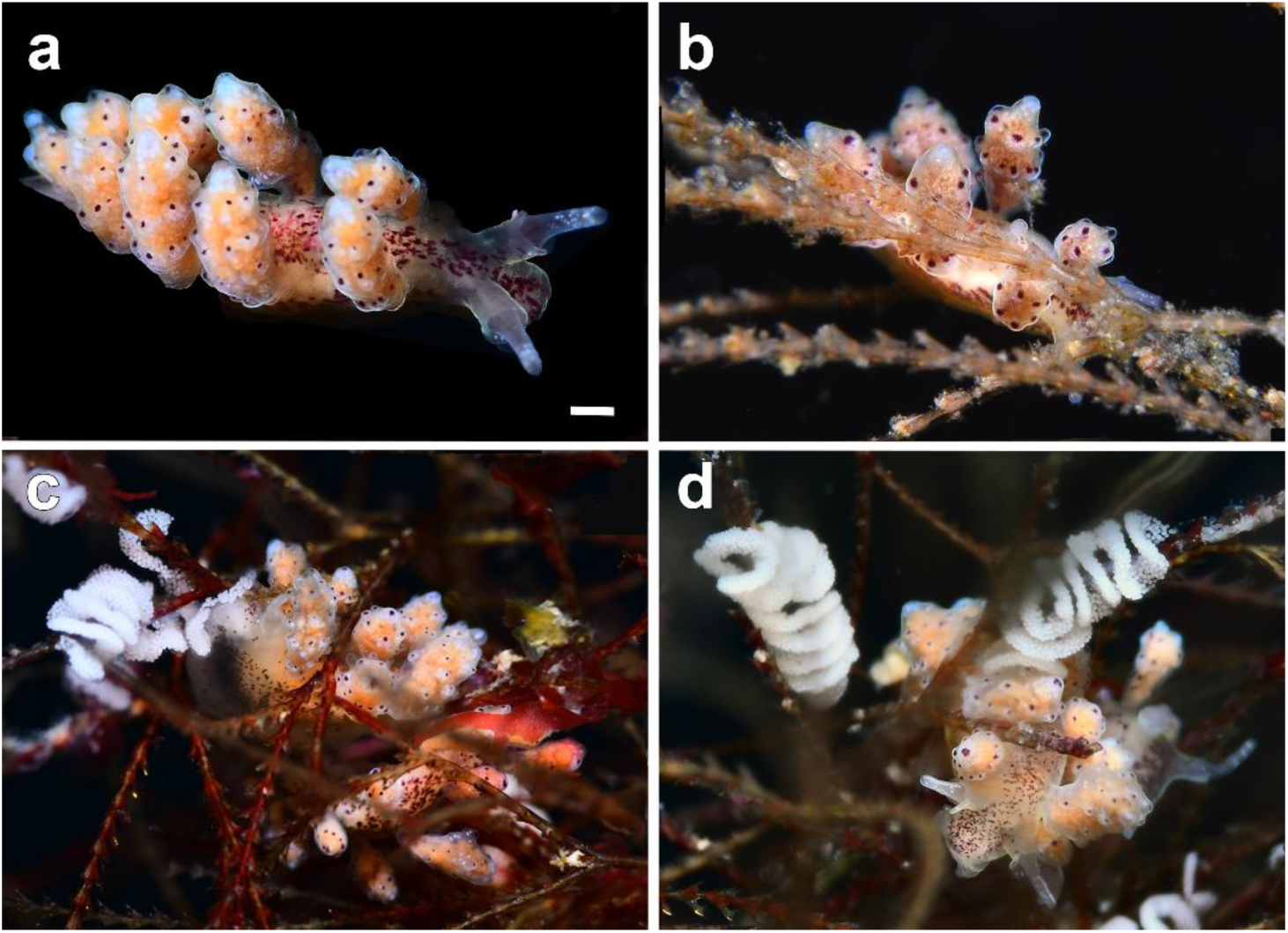
Photographs of live *Doto eireana* specimens **a** *D. eireana* (MCZ:Mala:406085) **b** same specimen on *Amphisbetia operculata* (Linnaeus, 1758) **e–d** *D. eireana* (MCZ:Mala:406086) with their egg masses on *Amphisbetia operculata*. Scale bar: 1 mm. Photos by Xavi Salvador

*Doto eireana* Lemche, 1976: 699–701; Ortea & Urgorri, 1978: 75; Trinchese, 1881: 92–93; Ballesteros, Madrenas & Pontes, 2016: 12; Picton & Morrow, 2023: 176–177.

**Material examined.** SPAIN · 3 spcs, L = 2–7 mm (one sequenced); Galicia, Viveiro, Playa de Area; 43°41’36.2“N, 7°34’34.9”W, 1.5 m depth; 01 Jul. 2022; X. Salvador leg.; MCZ:Mala:406084 · 2 spcs, L = 2–7 mm (one sequenced); same collecting place and date as for preceding,; X. Salvador leg.; MCZ:Mala:406086 · 1 spc., L = 5 mm (sequenced); same collecting place as for preceding, 2 m depth; 02 Jul. 2022; X. Salvador leg.; MCZ:Mala:406085.

**External morphology.** (Fig. 10, Fig. 5d) Body elongated, narrow, translucent, white background, with red to maroon irregular dots evenly distributed. Rhinophores white pigmented in apical half; rhinophoral sheath cylindrical, symmetrical, with red to maroon dots unevenly distributed on internal side. Dorsolateral appendages in 3–5 pairs, bulbous, internal part of base with red irregular dots; tubercles displayed in 3–5 crowns, blunt, widest just below tip, with apical dark maroon dots, not always present on apical tubercle.

**Egg mass.** (Fig. 10c–d, Fig. 5d) White, with a sinusoidal cord shape, 7–10 folds, arranged in linear disposition. Sometimes circularly wound to the hydrozoans.

**Ecology.** The specimens were found at shallow depths in vertical walls with an abundance of hydrozoans, especially *Amphisbetia operculata*, the same as *D. coronata* and *D. pinnatifida* (Martinsson et al., 2021).

**Life cycle.** Adults and egg masses are present from May to September. Mating behaviour was recorded between May to July based on different dive points on the Galician coast.

**Distribution.** North Eastern Atlantic coast (Lemche, 1976; OBIS, 2024); Iberian Peninsula (Ortea & Urgorri, 1978; Calado et al., 2003). The specimens cited in Catalonia (Ballesteros et al., 2016) and the French Mediterranean coast (GROC, 2024) seem misidentifications of *D. eireana*. Their pictures seem to correspond to *D. maculata*. The strict connection with the hydrozoan *Amphisbetia operculata* and its absence in these areas reinforces the fact that the species is not present in the Mediterranean.

**Remarks.** This species was described in the British Islands (Lemche, 1976) and found on the Atlantic side of the Iberian Peninsula (Ortea & Urgorri, 1978). Currently, there is no study providing molecular data to support the presence of *D. eireana* in the Mediterranean. The records of *D. eireana* in the Mediterranean (Ballesteros et al., 2016; GROC, 2024), given the description and images provided, most likely correspond to *D. maculata*.

Some differences are present between the two species. Our specimens of *D. eireana* present a redly pigmented and isodiametric rhinophoral sheath, being white and anteriorly extended in *D. maculata*. The base of the dorsolateral appendages is pigmented with irregular red spots and presents blunt tubercles in *D. eireana*, while they are unpigmented in *D. maculata* and the tubercles are elongated. Moreover, the oral veil has white spots only in *D. maculata*.

**Doto floridicola Simroth, 1888**

(Fig. 11a–c, Fig. 8c)

**Fig. 11.**
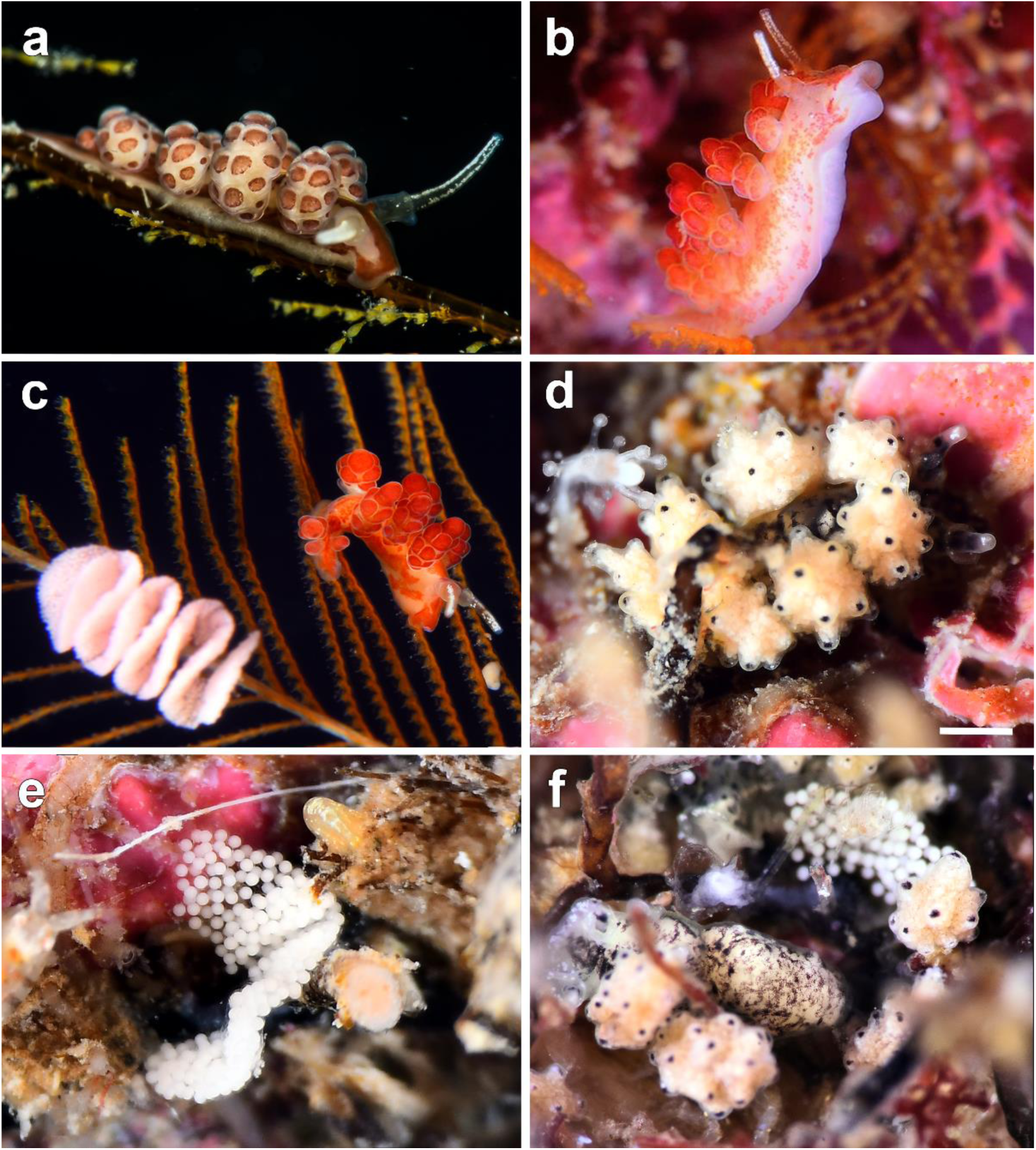
Photographs of live *Doto* species **a** *D. floridicola* (MCZ:Mala:393368) on *Aglaophenia elongata* **b** sequenced specimen of *D. floridicola* (MCZ:Mala:393367) on *Aglaophenia tubulifera* **c** *D. floridicola* with their egg mass on *Aglaophenia kirchenpaueri* **d–e** *D. fluctifraga* (MCZ:Mala:406087) and their egg mass **f** *D. fluctifraga* with their egg mass. Scale bar: 0.5 mm. Photos by Xavi Salvador, image of (a) courtesy by Joan Fernàndez

*Doto floridicola* Simroth, 1888: 219–222; Schmekel & Portmann, 1982: 164–166; Thompson, Cattaneo & Wong, 1990: 396; Ortea, Caballer & Moro, 2003: 181–187; Ortea et al., 2010: 83; Ortea & Moro, 2017: 6; Lombardo, 2021: 916; Picton & Morrow, 2023: 170–171.

*Doto susanae* Fez, 1962: 105–112.

**Material examined.** SPAIN · 1 spc. (sequenced); Catalonia, Girona, l’Escala, Cala Montgó; 42°06’28.7“N, 03°10’15.1”E, 1.5 m depth; 26 Dec. 2017; X. Salvador leg.; MCZ:Mala:393367 · 1 spc. (sequenced); Catalonia, Girona, l’Escala, Punta del Romaní; 42°06’53.1“N, 03°10’06.6”E, 12 m depth; 11 Aug. 2018; Joan Fernández leg.; MCZ:Mala:393368.

**External morphology.** (Fig. 11a–c, Fig. 8c) Body elongated, narrow, translucent, white background, with red or brown irregular spots that fuse forming longitudinal stripes, on dorsum and laterals, running from oral veil to tail. Rhinophores with a thin line of white dots on anterior and posterior part; rhinophoral sheath short, symmetrical, without dots. Dorsolateral appendages arranged in 4–6 pairs; tubercles displayed in 2–3 crowns, completely red, spaces between translucent.

**Egg mass.** (Fig. 11c, Fig. 8c) Light white to pink, arranged in a sinusoidal cord shape when deposited over the hydrozoan colonies.

**Ecology.** The specimens were found on *Aglaophenia tubulifera* (Hincks, 1861) and *A. kirchenpaueri* (Heller, 1868) on vertical walls exposed to the current and *A. elongata* Meneghini, 1845 in sandy patches. The recorded specimens were found active during the day.

**Life cycle.** Adults and egg masses are present all year. Mating behaviour was recorded between March and August (based on 138 records, 58 sampling dates, and 17 different localities).

**Distribution.** Atlantic Ocean: Azores Islands (Simroth, 1888), Galicia (Urgorri et al., 2011), Canary Islands (Ortea et al., 2003), Cabo Verde (Ortea & Moro, 2017), southwest of Britain and Ireland (Picton & Morrow, 2023); Gibraltar Strait (Garcia, 1983). Mediterranean: Cabo de Palos (Templado, 1982); eastern France (Wirz– Mangold & Wyss, 1958); Greece (Thompson et al., 1990); Italy (Thompson et al., 1990): Naples (Schmekel & Kress, 1977), Catania (Lombardo, 2021); Maltese Islands (Sammut & Perrone, 1998); Balearic Islands (Cervera et al., 2004); Mediterranean Spanish coast (Wollscheid Lengeling et al., 2001).

**Remarks.** This species is closely related to *D. lemchei*, both exhibit a similar shape and size (Picton & Morrow, 2023). *Doto floridicola* is distinguished from *D. lemchei* and other *Doto* species by its entirely coloured tubercles of red to brown colouration. Moreover, the distribution of both species overlaps in the Atlantic Ocean, specifically in northwest of Spain (Galicia), and in the southwest of Britain and Ireland (Picton & Morrow, 2023).

***Doto fluctifraga* Ortea & Perez, 1982**

(Fig. 11d–f, Fig. 8h)

*Doto fluctifraga* Ortea & Pérez, 1982: 79–83.

**Material examined.** SPAIN · 3 spcs, L = 2.5 mm (one sequenced); Canary Islands, Gran Canaria, Playa Amadores; 27°47’22.6“N, 15°43’30.0”W, 0.1 m depth; 07 Dec. 2021; X. Salvador leg.; MCZ:Mala:406087.

**External morphology.** (Fig. 11d, f, Fig. 8h) Body elongated, narrow, translucent, white background, with black irregular dots scattered on surface, densely interconnected. Rhinophores black pigmented, with white tip; rhinophoral sheath short, thin, symmetrical, with white dots on edge. Dorsolateral appendages arranged in 4 pairs, tubercles displayed in 2–3 crowns, rounded, with black dots on tip.

**Egg mass.** (Fig. 11e–f, Fig. 8h) Saccular, short, elongated and with a slight undulating shape; translucent in colour, with white eggs.

**Ecology.** The specimens were found in a floating line of buoys, over the stolons of *Pennaria disticha*. They were feeding and laying the egg masses at the base of the colony.

**Life cycle.** The cycle of this species is little known, with only a few records and, based on these records, probably present all year.

**Distribution.** Canary Islands (Ortea & Pérez, 1982); Azores (Ortea et al., 2010); Bahamas and Atlantic coast of North America (GBIF, 2024).

***Remarks*.** As for *D. fluctifraga*, this species can be easily distinguished from other *Doto* species by the black pigmentation on the rhinophores. There are no apparent morphological differences between *D. fluctifraga* and *D. cervicenigra*, although they are confirmed as two different species according to the SDT. They can be distinguished by their distribution and feeding preferences, as previously discussed above (see *D. cervicenigra*’s description). In Canarian waters, the type locality of *D. fluctifraga*, *D. eo* also presents black pigmentation on the rhinophores. *Doto eo* differs from *D. fluctifraga* by the salmon colour of the dorsolateral appendages and the conical and less prominent shape of their tubercles (Ortea & Moro, 2014).

***Doto fragaria* Ortea & Bouchet, 1989**

(Fig. 12, Fig. 8a)

**Fig. 12.**
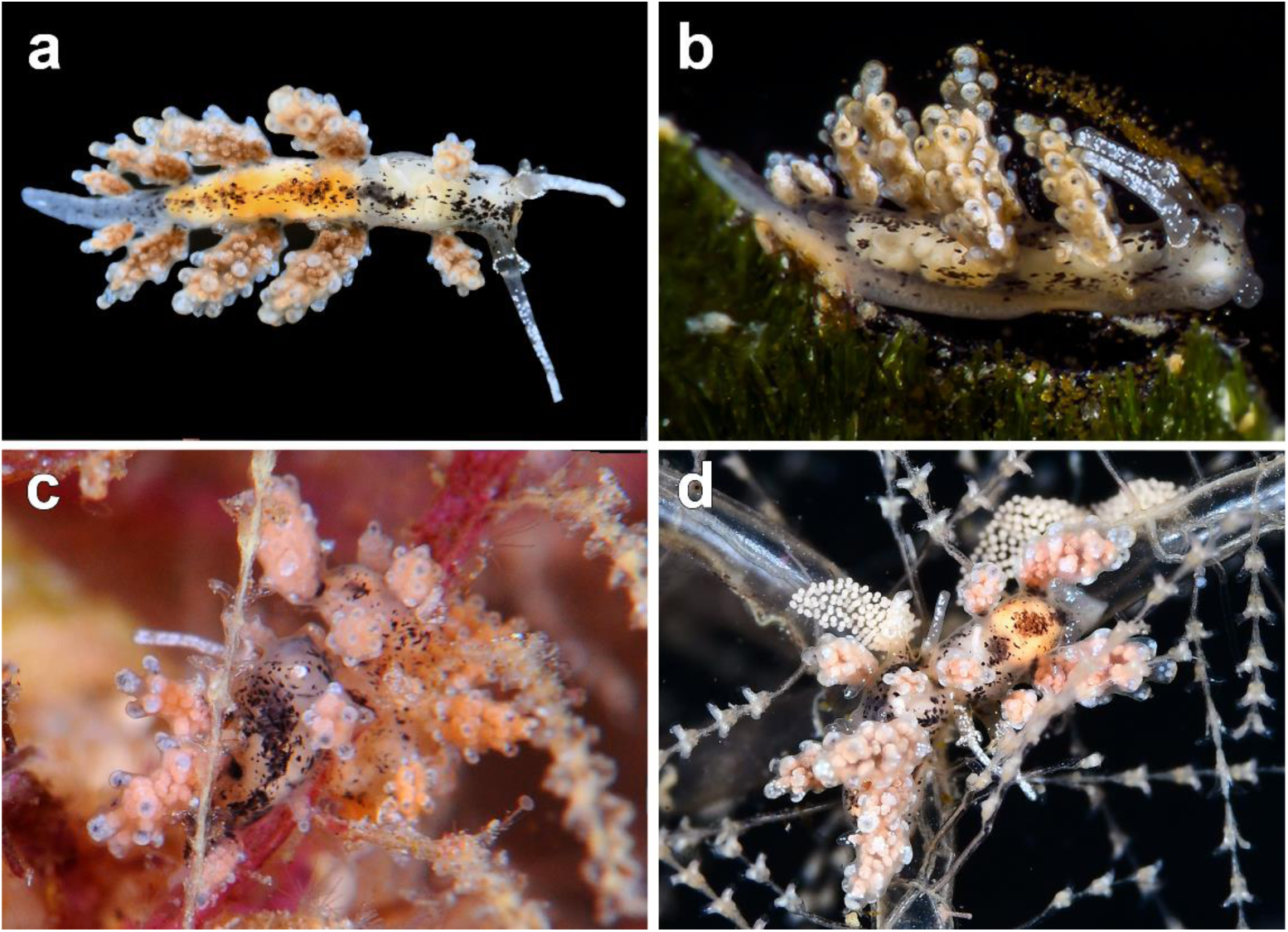
Photographs of live *Doto fragaria* specimens **a** *D. fragaria* **b** sequenced specimen (MCZ:Mala:393387) **c** specimens on *Dynamena* sp. **d** specimens with their egg mass on *Dynamena* sp. Photos by Xavi Salvador, image of (b) courtesy by Joan Fernàndez

*Doto fragaria* Ortea & Bouchet, 1989: 262–264.

**Material examined.** SPAIN · 1 spc. (sequenced); Catalonia, Girona, l’Escala, Punta del Romaní; 42°6’53.1“N, 3°10’6.6”E, 2 m depth; 11 Aug. 2018, Joan Fernàndez leg.; MCZ:Mala:393387.

**External morphology.** (Fig. 12, Fig. 8a) Body elongated, narrow, translucent, white to cream background, with black irregular markings scattered on surface. Rhinophores white pigmented; rhinophoral sheath short, extended anteriorly, with white dots on edge. Dorsolateral appendages arranged in 4–5 pairs, of strawberry colour, base black pigmented in internal part; tubercles displayed in 3–5 crowns, rounded, white in color, withcentral internal black dot.

**Egg mass**. (Fig. 12d, Fig. 8a) Translucent with tiny white eggs, arranged in a lineal disposition.

**Ecology.** The specimens were found at shallow depths, with an abundance of small hydrozoans colonies, especially the genera *Sertularia* and *Dynamena* Lamouroux, 1812 [probably *Sertularia distans*]. The specimens were active during the day.

**Life cycle.** Adults were found between January and February and from August to September; mating behaviour and egg masses were only recorded in September (based on 16 records, 7 sampling dates, and 4 different localities).

**Distribution.** Corsica (Ortea & Bouchet, 1989); Italy (Furfaro et al., 2020); Catalonia (GROC, 2024).

**Remarks.** The type locality of this species is Corsica, and it has been scarcely recorded afterwards. This species is very similar to *D. rosea*, but it is characterized by having longer, isodiametric dorsolateral appendages, and very rounded tubercles, with a black inner center (Ortea & Bouchet, 1989). Likewise, the egg masses are completely different, in *D. fragaria* they are smaller and saccular, with white tiny eggs, while in *D. rosea* the egg masses are long and sometimes lightly pigmented, arranged in a “S” shape disposition or circular.

**Doto koenneckeri Lemche, 1976**

(Fig. 13, Fig. 8e)

**Fig. 13.**
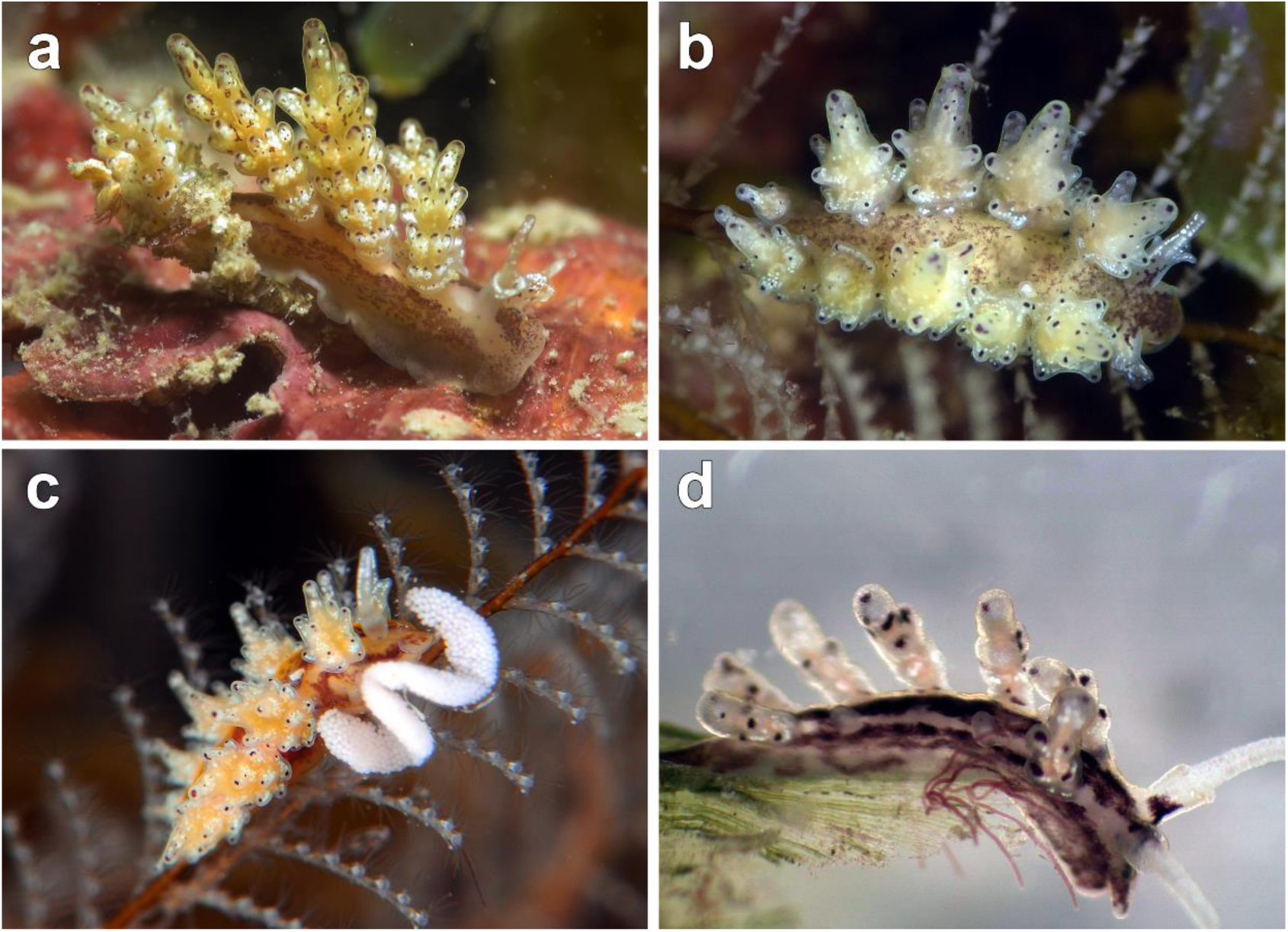
Photographs of live *Doto koenneckeri* specimens. Sequenced specimens **a** MCZ:Mala:393372 and **b** MCZ:Mala:393374 **c** specimen with their egg mass on *Aglaophenia pluma* (Linnaeus, 1758) **d** sequenced juvenile specimen (MCZ:Mala:393374). Photos by Xavi Salvador

*Doto koenneckeri* Lemche, 1976: 702–703; Ortea et al., 2010: 83–84; Moro, Ortea & Bacallado, 2016: 30–31; Picton & Morrow, 2023: 170–171.

**Material examined.** SPAIN · 1 spc. (sequenced); Catalonia, Girona, Tossa de Mar, Mar Menuda; 41°43’14.1“N, 2°56’26.1”E, 12 m depth; 03 Sep. 2016; Guillem Mas leg.; MCZ:Mala:393377 · 1 spc. (sequenced); Catalonia, Girona, Begur, Cala d’Aiguafreda; 41°57’51“N, 3°13’39.8”E, 6 m depth; 04 Jan. 2018; X. Salvador leg.; MCZ:Mala:393370 · 1 spc. (sequenced); same collecting place and date as for preceding, 6 m depth; Robert Fernández-Vilert leg.; MCZ:Mala:393373 · 1 spc. (sequenced); same collecting place as for preceding, 6 m depth; 09 Apr. 2018; Robert Fernández-Vilert leg.; MCZ:Mala:393374 · 3 spcs (one sequenced); Catalonia, Girona, l’Escala, Punta del Romaní; 42°6’53.1“N, 3°10’6.6”E, 12 m depth; 29 Mar. 2018; Robert Fernández-Vilert leg.; MCZ:Mala:393372.

**External morphology.** (Fig. 13, Fig. 8e) Body elongated, narrow, translucent, white to cream background, with brown irregular spots, forming longitudinal stripes, on dorsum and laterals, running from oral veil to tail.

Rhinophores white pigmented; rhinophoral sheath extended anteriorly, white markings on edge, base brown pigmented. Dorsolateral appendages arranged in 5–7 pairs, last pair smaller; tubercles displayed in 3–6 crowns, fusiform, elongated, apical one protruding, with one brown dot in ‘comma-shaped’ in apex of each tubercle, irregular brown dots between them. Juvenile specimens lacking white pigmentation on rinhophores and rhinophoral sheath, anterior extension of rhinophoral sheath absent, tubercles not fully elongated.

**Egg mass.** (Fig. 13c, Fig. 8e) White, displayed in a sinusoidal cord shape arranged over the center of hydrozoan colonies.

**Ecology.** The specimens were found at shallow depths, usually well illuminated, with the presence of *Aglaophenia pluma*. The specimens were found more active during the day when they lay their egg masses at the base or hide in the colony.

**Life cycle.** Adults present all over the year, especially abundant between February and June; egg masses are recorded all over the year; mating behaviour is only recorded between February and June (based on 581 records, 221 sampling dates, and 35 different localities).

**Distribution.** British Islands (Lemche, 1976; OBIS, 2024); Norway (Lemche, 1976); Portugal (Cervera et al., 2004); Azores Islands (Calado, 2002); Canary Islands (Moro et al., 2016); Spain: Galicia, Cantabria Sea, Alboran Sea, Catalonia coast (Cervera et al., 2004), Mallorca (GROC, 2024), Ceuta (Ortea et al., 2010); French Mediterranean coast (GBIF, 2024); Italy: Legurian coast (Betti et al., 2015), Sardinia coast (Trainito & Doneddu, 2015).

**Remarks.** This species resembles the closely related species *D. paulinae* but presents a white pigmentation on the rhinophoral sheath. The tubercles of the dorsolateral appendages are elongated with brown comma-shaped spots, which are entirely brown, except for the apical one, in *D. paulinae*. Moreover, the lateral pigmented bands of *D. koenneckeri* are absent in *D. paulinae*. The elongated apical tubercle and the comma-shaped markings on the tubercles are diagnostic characteristics of *D. koenneckeri* (Lemche, 1976). In this study, the specimens fixed in 96% EtOH had lost pigmentation and swollen dorsal appendages, thus making them unrecognizable among other species, highlighting the importance of life pictures for the correct identification of the specimens.

Phylogenetic analyses based on ML support the validity of *D. koenneckeri* and *D. paulinae*, whereas BI analyses lack sufficient support for this relationship. PTP analysis distinguishes both species, while ASAP clusters them together. Furthermore, these results suggest that specimens identified as *D. koenneckeri* from the Mediterranean and the Azores, together with an unidentified species, were misidentified and correspond to *D. paulinae*.

Similarly, specimens of *D. dunnei* that were grouped with *D. koenneckeri* also suggest possible misidentifications. Despite these inconsistencies, we maintain both species to be valid based on ML, PTP, and morphological differences.

***Doto maculata* (Montagu, 1804)**

(Fig. 14, Fig. 5e)

**Fig. 14.**
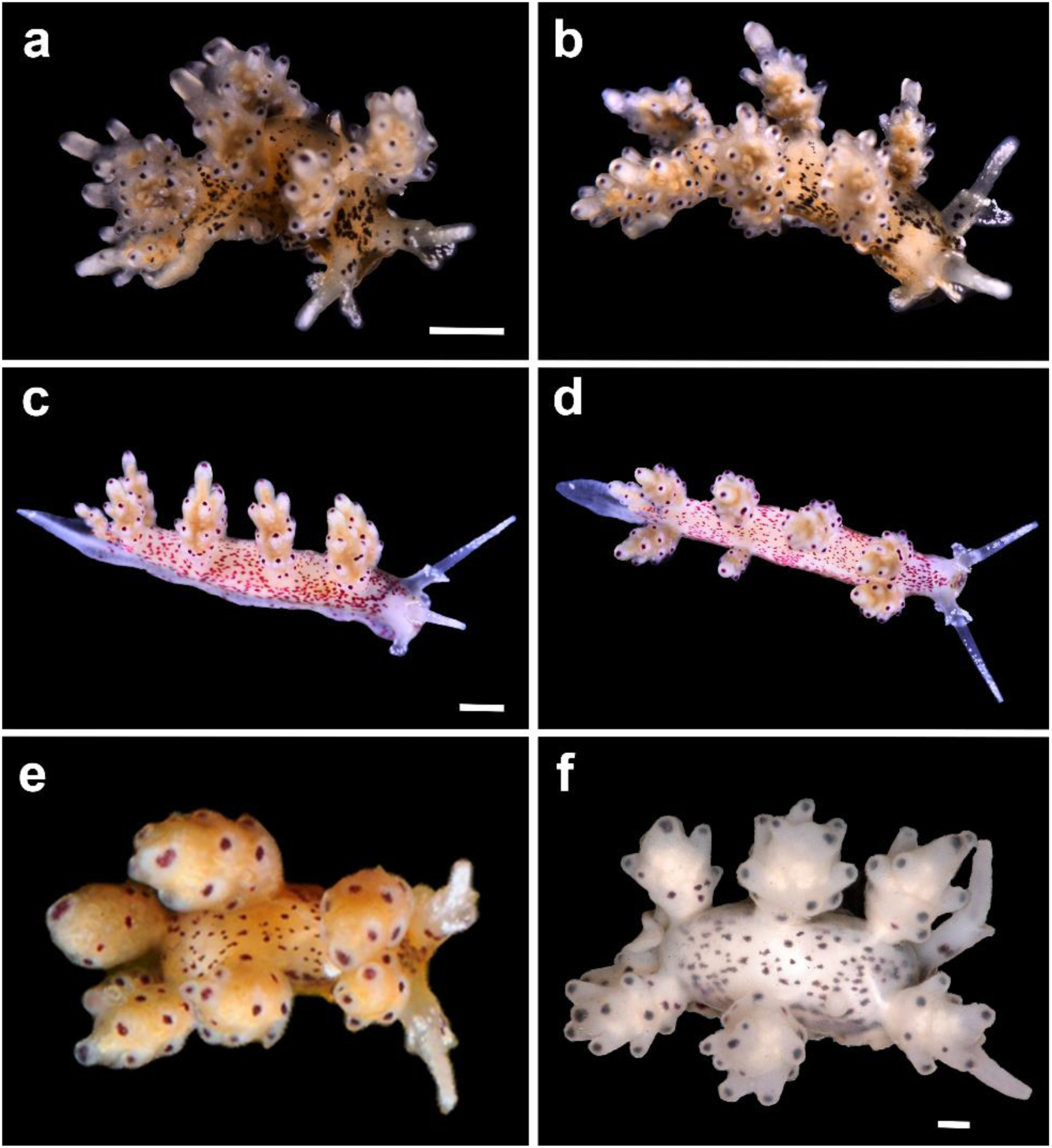
Photographs of live *Doto maculata* specimens, showing colouration variation of the Mediterranean taxa studied **a–b** specimen (MCZ:Mala:406088) in top and side view **c–d** specimen (MCZ:Mala:406089) in lateral and dorsal views **e–f** specimen MCZ:Mala:393361, (e) live and (f) preserved. Photos by Xavi Salvador, image of (f) courtesy by Juan Moles

*Doris maculata* Montagu, 1804: 61–85.

*Doto maculata*: Lemche, 1976: 696–697; Lombardo & Marletta, 2020: 916–917; Picton & Morrow, 2023: 192–193.

**Material examined.** SPAIN · 1 spc., L = 2.5 mm (sequenced); Catalonia, Girona, Sant Feliu de Guíxols, Cala Maset caves; 41°47’10.5“N, 3°2’44.6”E, 1.3 m depth; 07 Jun. 2020; X. Salvador leg.; MCZ:Mala:406088 · 1 spc., L = 5 mm (sequenced); same collecting place as for preceding, 2 m depth; 15 May 2020; X. Salvador leg.; MCZ:Mala:406089 · 1 spc., L = 5 mm (sequenced); same collecting place as for preceding, 0.1 m depth; 07 May 2018; X. Salvador leg.; MCZ:Mala:393361.

**External morphology.** (Fig. 14, Fig. 5e) Body elongated, narrow, translucent, cream background, with black brownish or red to pink irregular dots scattered all over body. Oral veil with white pigmentation on margin to base of rhinophores. Rhinophore apical half white; rhinophoral sheath projected anteriorly, with white dots on edge, sometimes with black or red dots on base. Dorsolateral appendages arranged in 3–5 pairs; tubercles displayed in 3–4 crowns, elongated, with several black spots between and within tubercles, especially in the lower part. Juvenile specimens present tubercles less elongated.

**Egg mass.** (Fig. 5e) White, displayed in a meandering cord shape.

**Ecology.** The specimens were found on vertical floating ropes in sand bottoms with high coverage of filamentous white to translucent hydrozoans and in shadow vertical walls with high coverage of unidentified hydrozoans. The slugs lay the egg masses over the hydrozoan colonies.

**Life cycle.** The specimens collected were found during spring, between May and June, with water temperatures over 20 °C. A 4th non-sequenced specimen was found in January, probably their life cycle in the Mediterranean Sea corresponds to the colder months (January to June).

**Distribution.** Northeast Atlantic (Lemche, 1976; GBIF, 2024); northwestern Spanish coast (Urgorri & Besteiro, 1986); Catalan coast (this study); Sicily (Lombardo & Marletta, 2020).

**Remarks.** Originally described from the South of the UK (Montagu, 1804), this species was considered a synonym of *D. coronata* until re-established its taxonomic validity (Lemche, 1976). Lemche (1976) states that there is no pigmentation on the rhinophoral sheath in their Atlantic specimens and that they have reddish or brownish spots at the apical part of each tubercle, usually except for the apical one. However, our Mediterranean specimens present spots on the inner part of the rhinophoral sheath, similar to the specimens examined by Urgorri & Besteiro (1986) from Galicia (Atlantic). Regarding the tubercles, all specimens of *D. maculata* examined have a red or brown spot on the apical tubercle. In contrast, in some of the examined specimens of *D. eireana*, the absence of pigmentation on the apical tubercle, a characteristic of *D. maculata* and not *D. eireana* according to Lemche (1976), can be observed. Because of this, there may be mistaken identifications between *D. eireana* and *D. maculata* (Ballesteros et al. 2016). The morphological differences between both species are detailed in the section of *D. eireana*. Overall, *D. maculata* specimens from the Mediterranean exhibit a white pigmentation on the rhinophores, the rhinophoral sheath, and their oral veil (Lombardo, 2021), and there might be more than one spot per tubercle, also between them. These characteristics are not mentioned in Lemche’s (1976) description. Another possible confusion, due to the long dorsolateral appendages and elongated tubercles is with *D. koenneckeri*, a species that presents comma-shaped spots in the tips instead to the usual dots of *D. maculata*.

In this study, juvenile specimens of *D. maculata* were observed with less elongated tubercles compared to adults, being somewhat more like *D. eireana*. Once the juveniles are fixed in 96% EtOH the dorsolateral appendages tend to extend compared to their natural state (Fig. 14e–f).

This species is reported in the northeastern Atlantic, from the British Islands to the northwest of Spain, and in the eastern Mediterranean. In this study, the species is reported for the first time on the Catalan coast (western Mediterranean).

According to the SDT, our *D. maculata* specimens from the Mediterranean belong to two putative species. This intraspecific divergence could be due to the variability and limited availability of the marker used for the SDTs (see Fig. S1). Future studies would need to increase the number of markers for this species to validate these results. However, despite the COI divergence, we maintain all the new Mediterranean specimens as the same species, supported by the absence of external morphological differences, but for the dot colouration.

**Doto millbayana Lemche, 1976**

(Fig. 15, Fig. 5c)

**Fig. 15.**
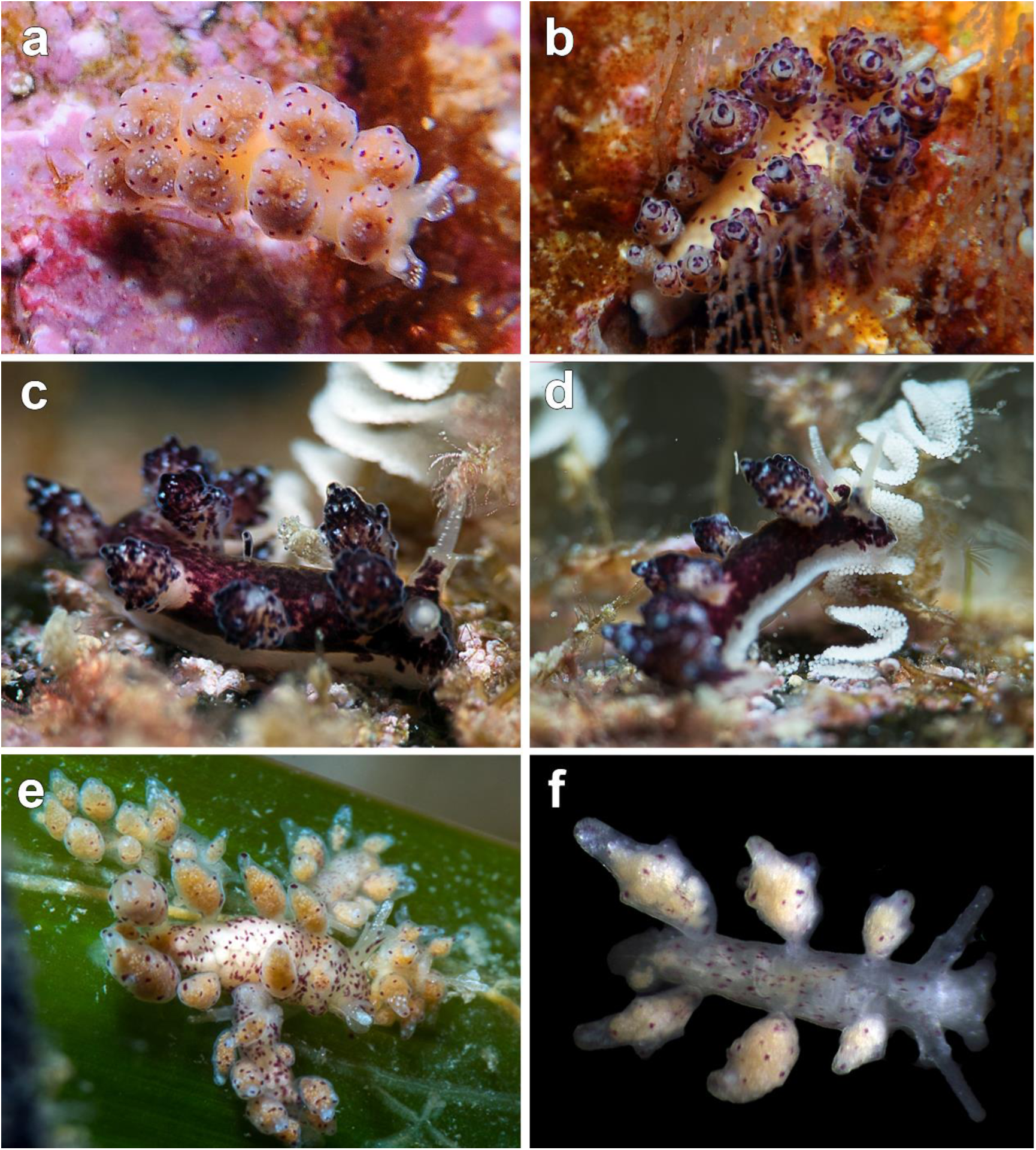
Underwater photographs of *Doto millbayana* specimens, showing morphological variation of the Mediterranean taxa studied **a** *Doto* cf. *coronata* (MCZ:Mala:393355) **b** *Doto* cf. *dunnei* (MCZ:Mala:393364) on rocky substrate **c–d** *Doto* ‘*dunnei*’ (MCZ:Mala:393365) with their egg mass on rocky substrate **e** *Doto* cf. *coronata* (MCZ:Mala:393357) on *Posidonia* meadows **f** *Doto* cf. *coronata* (MCZ:Mala:393363) found living on *Posidonia* meadows. Photos by Xavi Salvador

*Doto millbayana*: Lemche, 1976: 701–702; Ortea & Urgorri, 1978: 76–78; Picton & Morrow, 2023: 194–195.

*Doto dunnei*: **new synonymy**.

**Material examined.** SPAIN · 1 spc. (sequenced); Catalonia, Girona, Sant Feliu de Guíxols, Cala Maset caves; 41°47’10.5“N, 3°2’44.6”E, 1 m depth; 14 Feb. 2018; X. Salvador leg.; MCZ:Mala:393364 · 2 spcs (one sequenced); same collecting place as for preceding, 0.1 depth; 19 Mar. 2018; X. Salvador leg.; MCZ:Mala:393362 · 1 spc. (sequenced); same collecting place as for preceding, 0.1 m depth; 09 Mar. 2018; X. Salvador leg.; MCZ:Mala:393360 · 1 spc. (sequenced); Catalonia, Girona, Begur; Cala d’Aiguafreda; 41°57’51“N, 3°13’39.8”E, 6 m depth; 04 Jan. 2018; Irene Figueroa leg.; MCZ:Mala:393354 · 1 spc. (sequenced); same collecting place as for preceding, 15 m depth; 07 Apr. 2018; Robert Fernández-Vilert leg.; MCZ:Mala:393365 · 1 spc. (sequenced); Catalonia, Girona, Begur, Sa Tuna; 41°57’36.9“N, 3°13’51.2”E, 3 m depth; 25 Jan. 2018; X. Salvador leg.; MCZ:Mala:393355 · 1 spc. (sequenced); Catalonia, Girona, l’Escala, Punta del Romaní; 42°6’53.1“N, 3°10’6.6”E, 8 m depth; 25 Feb. 2018; Robert Fernández-Vilert leg.; MCZ:Mala:393357 · 1 spc. (sequenced); collecting place and date as for preceding, 8 m depth; Robert Fernández-Vilert leg.; MCZ:Mala:393363 · 2 spcs (one sequenced); Catalonia, Girona; Blanes, Punta Santa Anna; 41°40’25.8“N, 2°48’6.2”E, 8 m depth; 04 Dec. 2018; Robert Fernández-Vilert leg.; MCZ:Mala:393366 · 1 spc. (sequenced); Catalonia, Girona, Tossa de Mar, Mar Menuda; 41°43’14.1“N, 2°56’26.1”E, 5 m depth; 18 Feb. 2018; Guillem Mas leg.; MCZ:Mala:393356.

**External morphology.** (Fig. 15, Fig. 5c) Body elongated, narrow, translucent, cream to white background, with purple or red to maroon irregular dots, in some specimens almost completely purple pigmented. Rhinophores white pigmented; rhinophoral sheath short, extended anteriorly, with white dots on edge, purple or maroon pigmented at base. Dorsolateral appendages arranged in 4–6 pairs; tubercles displayed in 2–4 crowns, barely defined, except apical one, with purple, maroon or red dots in each tubercle, and discontinuous dot circles around them. Specimens showing purple colouration with well-defined tubercle tips.

**Egg mass.** (Fig. 15c–d, Fig. 5c) White, displayed in a meandering cord shape. Depending on the substrate it can be either linear, with up to 6 folds, or deposited in a circle shape.

**Ecology.** The specimens found on rocky substrates on *Kirchenpaueria halecioides* (Alder, 1859) or *K. pinnata* (Linnaeus, 1758). Populations of *D. millbayana* found on *Posidonia oceanica* (Linnaeus) Delile, 1813 meadows are smaller in size and feed on *Tridentata perpusilla* Stechow, 1919 and *Obelia* sp., where they lay the egg masses in circles. The specimens were found active during daylight.

**Life cycle.** Adults are present between January and June; mating behaviour and egg masses are recorded between February and May (based on 283 records, 81 sampling dares, and 9 different localities). Nevertheless, due to early misidentifications between *D. dunnei* and *D. coronata*, the life cycle could probably be more extensive.

**Distribution.** British Islands (Lemche, 1976; OBIS, 2024); Portugal (Cervera et al., 2004); Spain: Asturias (Ortea & Urgorri, 1978), Galicia, Strait of Gibraltar (Cervera et al., 2004), Catalonia (cited as *D. dunnei* and *D. coronata* in Ballesteros et al, 2016; this study).

**Remarks.** In the original description of *D. millbayana* from South UK the presence of an irregular distribution of spots on dorsolateral appendages and tubercles was considered diagnostic (Lemche, 1976), a feature lacking in *D. coronata*. All our specimens from the Mediterranean present this diagnostic feature but show a great chromatic and morphological variation. The specimens found in *Posidonia* meadows (traditionally attributed to *D. coronata*; Fig. 15a, e–f and Fig. 5c2) have smoother dorsolateral appendages, without profusely marked tubercles except for the apical one, and present an irregular distribution of maroon markings between tubercles. In contrast, the specimens found on rocky substrates (traditionally attributed to *D. dunnei*; Fig. 15b–d and Fig. 5c3) have dorsolateral appendages with rounded tubercles, presenting purple dots in each tubercle, with discontinuous dot circles around, and the purple pigmented body, in some specimens almost completely.

Furthermore, these two morphotypes also differ in diet, as mentioned in the section on ecology. We found morphological variability in specimens of *D. millbayana* from different substrates, but all material examined from the Mediterranean is characterized by the presence of more than one dot per tubercle in the dorsolateral appendages. In this study, molecular data support the presence of *D. millbayana* in the northwest Mediterranean.

Here, molecular data and SDT analyses show that *D. millbayana* and *D. dunnei* are the same species, as in previous molecular studies (Shipman & Gosliner, 2015). Martinsson et al. (2021) suggested that *D. millbayana* and *D. dunnei* should remain separate species, as they mention that they can be differentiated by minor radular characters, feeding habits, the shape of pseudobranchs, and the distribution and quantity of body spots.

Molecular data and the extensive morphological variation of *D. millbayana* from the Atlantic to the Mediterranean support the junior synonymy of *D. dunnei* to *D. millbayana*. Morphological differences are here attributed to intraspecific variation among *D. millbayana* populations.

***Doto paulinae* Trinchese, 1881**

(Fig. 16, Fig. 8d)

**Fig. 16.**
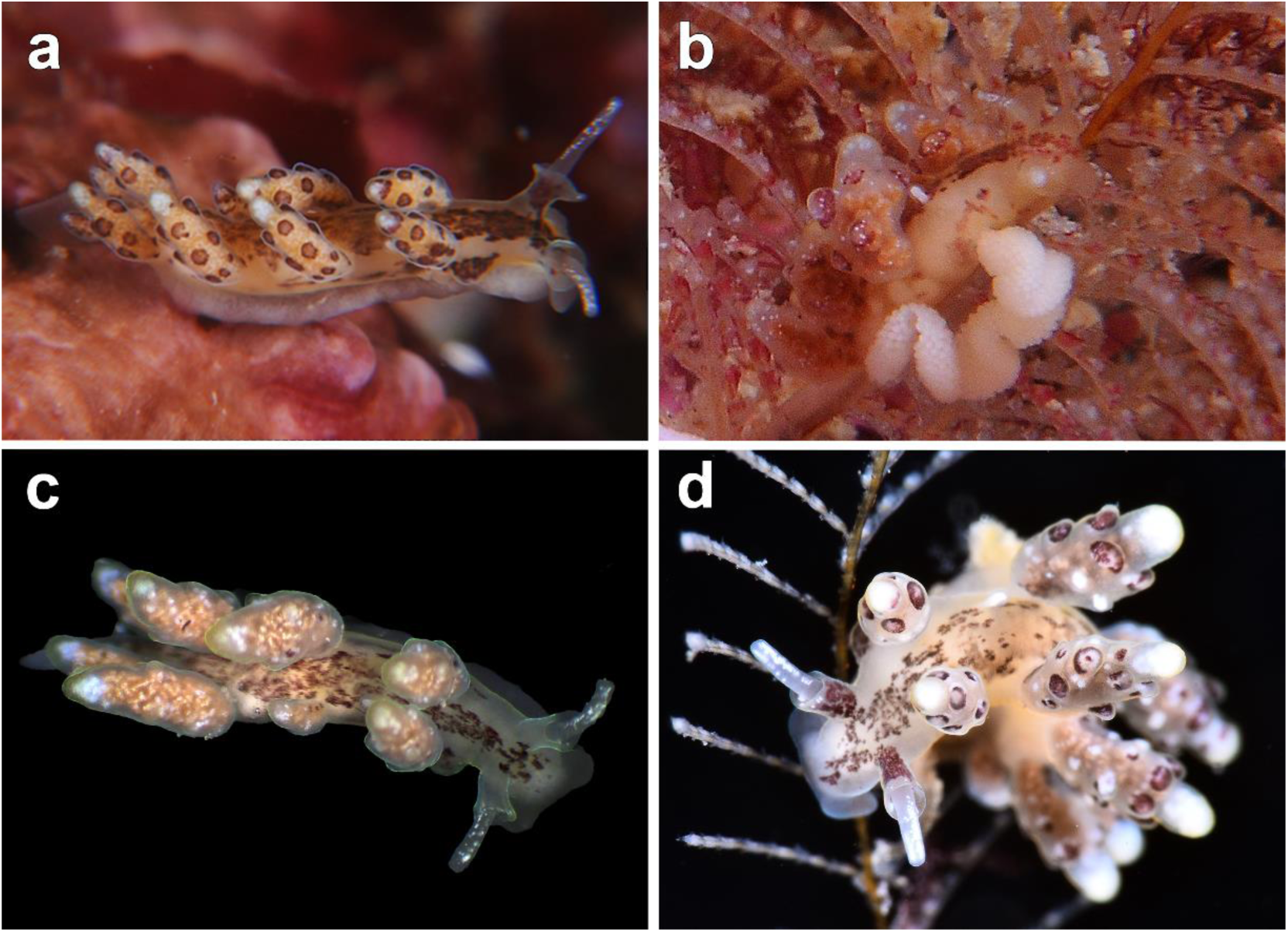
Photographs of live *Doto paulinae* specimens **a** sequenced specimen MCZ:Mala:393383 **b** sequenced specimen MCZ:Mala:393379 with their egg mass on *Aglaophenia* sp. **c** sequenced juvenile specimen MCZ:Mala:393382 **d** specimen on *Aglaophenia* sp. Photos by Xavi Salvador

*Doto paulinae* Trinchese, 1881: 93; Schmekel & Portmann, 1982: 166–167.

*Doto styligera* Hesse, 1872: 345–348.

**Material examined.** SPAIN · 1 spc. (sequenced); Catalonia, Girona, l’Escala, Punta del Romaní; 42°6’53.1“N, 3°10’6.6”E, 5 m depth; 24 Jan. 2018; X. Salvador leg.; MCZ:Mala:393379 · 1 spc. (sequenced); same collecting place as for preceding, 6 m depth; 25 Feb. 2018; Guillem Mas leg.; MCZ:Mala:393382 · 1 spc. (sequenced); same collecting place and date as for preceding, 6 m depth; Guillem Mas leg.; MCZ:Mala:393384 · 1 spc. (sequenced); same collecting place as for preceding, 1 m depth; 24 Apr. 2018; X. Salvador leg.; MCZ:Mala:39338 · 1 spc. (sequenced); same collecting place as for preceding, 5 m depth; 04 Mar. 2016; Guillem Mas leg.; MCZ:Mala:393375.

**External morphology.** (Fig. 16, Fig. 8d) Body elongated, narrow, translucent, white to cream background, with brown irregular spots that form a longitudinal stripe on the dorsum. Rhinophores white pigmented; rhinophoral sheath short, extended anteriorly, base brown pigmented. Dorsolateral appendages displayed in 3–5 pairs; tubercles arranged in 2–3 crowns, rounded, brown, except apical tubercle white and cupular. Juvenile specimens lack brown pigmentation on rhinophoral sheath, tubercles shorter and white.

**Egg mass.** (Fig. 16b, Fig. 8d) White, displayed in a sinusoidal cord, extremely short (from two to three loops), lineally deposited over hydrozoan colonies (normally out of light exposure).

**Ecology.** The specimens were found at shallow depths, inside caves or in shadowy sloping walls with moderate hydrodynamics and abundance of *Aglaophenia* sp. The specimens are more active during the night, being at the base of the hydrozoan colony during the day.

**Life cycle.** Adults are found during January and September; egg masses are recorded during all the life cycle; mating behavior is only recorded in February (based on 39 records, 32 sampling dates, and 9 different localities).

**Distribution.** Italy (Trinchese, 1881): Sardinia (Trainito et al., 2015); NE Spain (Cervera et al., 2004); Catalan coast (Ballesteros, 2007); French Mediterranean coast (GBIF, 2024).

**Remarks.** This species is characterized by the presence of brown, completely pigmented tubercles, except for the apical one, which is cupular and white. *Doto paulinae* resembles its more closely related species *D. koenneckeri* (see the description above). Phylogenetic analyses based on ML and PTP analyses conducted, along with differences in external morphology, here support the validity of these two species.

**Doto pygmaea Bergh, 1871**

(Fig. 17, Fig. 8f)

**Fig. 17.**
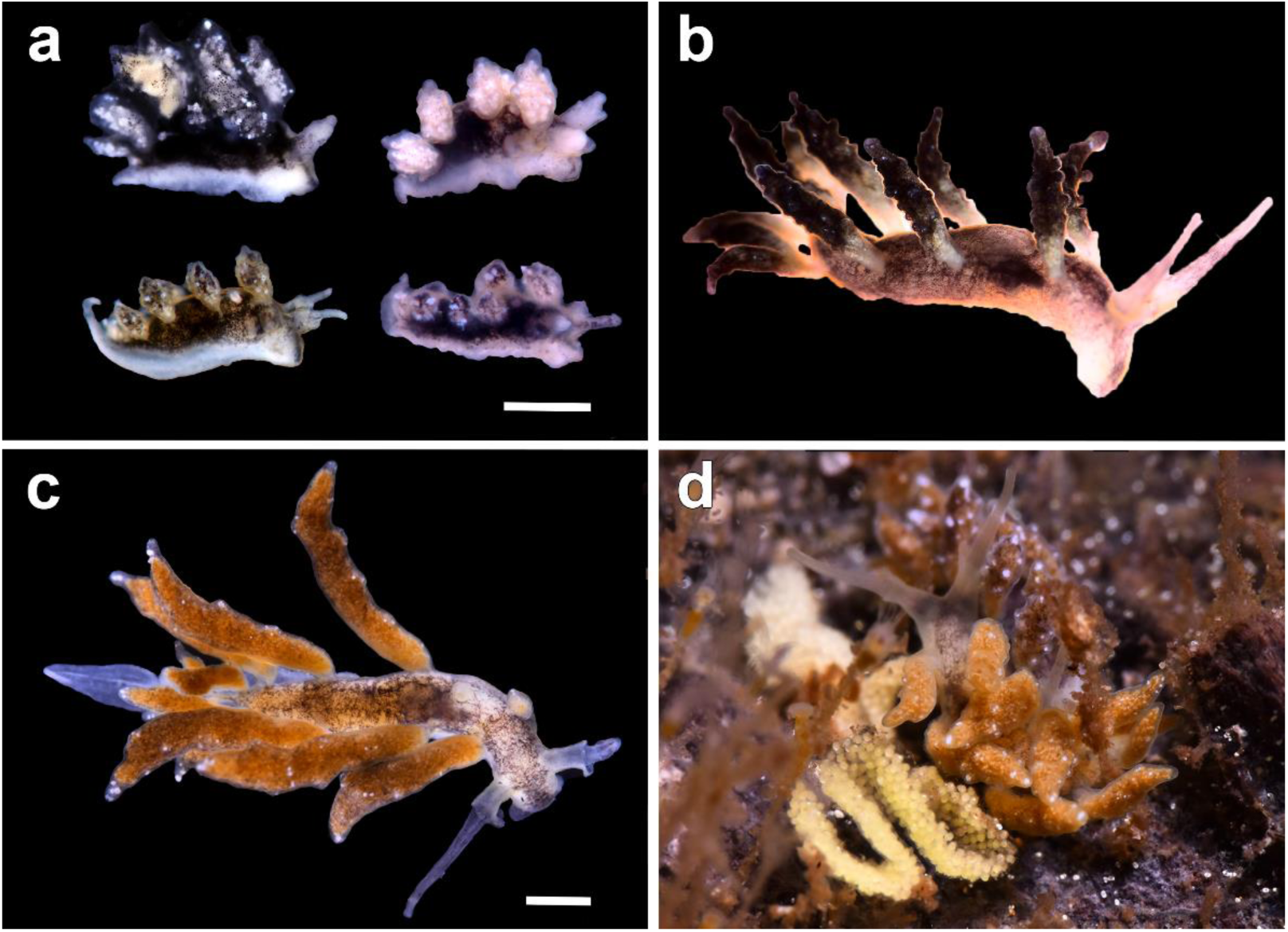
Photographs of live *Doto pygmaea* specimens **a** sequenced specimen MCZ406091 found living on floating beach buoys **b** specimen found on buoys **c–d** sequenced specimen MCZ:Mala:406090 (d) on floating branch with their yellow egg mass. Scale bar: 0.5 mm. Photos by Xavi Salvador

*Doto pygmaea* Bergh, 1871: 1277–1280; Ortea, Moro & Espinosa, 1997: 126–128; Sanvicente-Añorve et al., 2012: 456; Ballesteros, Madrenas & Pontes, 2016: 12–14; Salvador, Fernández-Vilert & Moles, 2022: 280–282.

*Doto doerga* Marcus & Marcus, 1963: 39–41; Schmekel & Portmann, 1982: 162–164.

**Material examined.** SPAIN · 9 spcs, L = 1–2 mm (one sequenced); Catalonia, Girona, Begur, Platja Fonda; 41°56’30“N, 3°13’03”E, 0 m depth; 22 May 2020; X. Salvador leg.; MCZ:Mala:406091 · 1 spc., L = 4 mm (sequenced); Catalonia, Girona, Sant Feliu de Guíxols, Cala Maset caves; 41°47’10.5“N, 3°2’44.6”E, 0 m depth; 01 Feb. 2019; X. Salvador leg.; MCZ:Mala:406090.

**External morphology.** (Fig. 17, Fig. 8f) Body elongated, narrow, white or beige background with black patches, except on tail, densely interconnected, up to completely black. Rhinophores translucent; rhinophoral sheath extended anteriorly in adults, black or sometimes depigmented. Dorsolateral appendages arranged in 4–5 pairs, internal side arched, (very) elongated, with variable color between black, beige and white; tubercles little protruded in external part, smooth internally, white pigmentation only in external part.

**Egg mass.** (Fig. 17d, Fig. 8f) White to yellow, displayed in a meandering cord shape. Depending on the substrate and the size of the specimens it can be either linear, with up to 6 folds, or deposited in a circle shape in adults in large floating objects.

**Ecology.** The specimens were found on floating objects, e.g., plastics, beach buoys, floating locks or branches with coverage of hydrozoans.

**Life cycle.** The species is present all over the year, specially during the summer months, with presence of floating signalling buoys in the beach, that cover by hydrozoans in 2–3 weeks. In larger floating objects, the specimens found were larger and with more elaborated egg masses than in the buoys or small plastic debris.

**Distribution.** Bahamas and Cuba (Sanvicente-Añorve et al., 2012); Sargasso Sea (Bergh,1871); Canary Islands (Ortea et al., 1997); western Mediterranean Sea (Schmekel & Portmann, 1982): Levantine coast (Cervera et al., 2004), Catalonia (Salvador, Fernández Vilert & Moles, 2022); Italy (Schmekel & Portmann, 1982).

**Remarks.** This species is easily distinguished from other conspecific *Doto* species by the asymmetric shape of its dorsolateral appendages, with a smooth internal side lacking tubercles and pseudobranchs (Bergh, 1871; Ortea et al., 1997). They are found on floating debris feeding on the hydrozoans *Aglaophenia pluma* (McDonald & Nybakken, 1997), *Clytia hemisphaerica* (Linnaeus, 1767) (Micaroni et al., 2018), and *Obelia geniculata* (Linnaeus, 1758) (Schmekel & Portmann, 1982).

***Doto rosea* Trinchese, 1881**

(Fig. 18a–d, Fig. 8b)

**Fig. 18.**
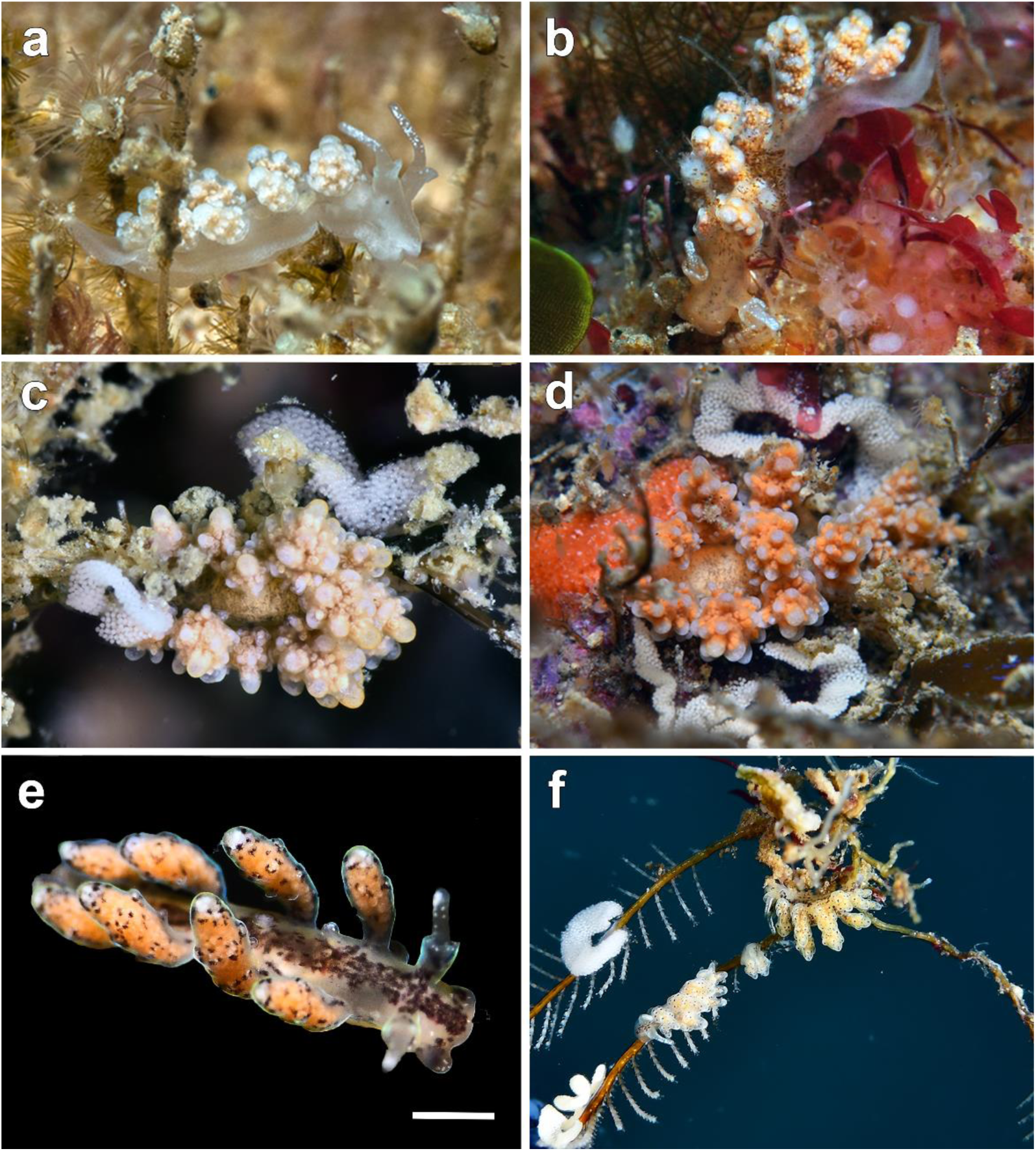
Photographs of live *Doto* species. Sequenced specimens of *D. rosea* **a** MCZ:Mala:393351 and **b** MCZ:Mala:393390 **c** *D. rosea* with their egg mass on *Eudendrium* sp. **d** *D. rosea* with their circular egg mass on rocky substrate **e** sequenced specimen of *D. verdicioi* (MCZ:Mala:406092) **f** *D. verdicioi* with their egg mass on hydrozoan. Scale bar: 0.2 mm. Photos by Xavi Salvador

*Doto rosea* Trinchese, 1881: 92; Schmekel & Portmann, 1982: 167–169; Thompson, Cattaneo & Wong, 1990:

398–400; Ballesteros, Madrenas & Pontes, 2016: 14.

*Doto aurea* Trinchese, 1881: 92.

*Doto cinerea* Trinchese, 1881: 92–93.

**Material examined.** SPAIN · 1 spc. (sequenced); Catalonia, Girona, Tossa de Mar, Mar Menuda; 41°43’14.1“N, 2°56’26.1”E, 6 m depth; 18 Feb. 2018; Irene Figueroa leg.; MCZ:Mala:393350 · 2 spcs (one sequenced); Catalonia, Girona, Sant Feliu de Guíxols; Cala Maset caves; 41°47’10.5“N, 3°2’44.6”E, 0.5 m depth; 2 Jan. 2018; X. Salvador leg.; MCZ:Mala:393390 · 1 spc. (sequenced); Catalonia, Girona, l’Escala, Punta del Romaní; 42°6’53.1“N, 3°10’6.6”E, 4 m depth; 25 Feb. 2018; Irene Figueroa leg.; MCZ:Mala:393351 · 2 spcs (one sequenced); Catalonia, Girona, Cadaqués, Cova de l’Infern; 42°19’02.9“N, 3°19 ‘12.1”E, 2 m depth; 18 Sep. 2018; X. Salvador leg.; MCZ:Mala:393369.

**External morphology.** (Fig. 18a–d, Fig. 8b) Body elongated, narrow, translucent, white to cream background, with tiny black marks. Rhinophores white pigmented; rhinophoral sheath, base unpigmented, with white dots on edge. Dorsolateral appendages displayed in 5–6 pairs, oval, elongated, background color variable between white, grey, brown, or pink; tubercles displayed in 3–4 crowns, rounded, apical part completely white.

**Egg mass.** (Fig. 18c–d, Fig. 8b) White, displayed in an undulated cord shape. Depending on the substrate it can be either linear (e.g., over *Eudendrium* Ehrenberg, 1834), with four undulations (not completely folded), or circular in rocky substrate.

**Ecology.** This species feeds on hydrozoans of the genera *Eudendrium*, *Sertularella*, or *Campanularia*. The specimens were found active during the day and crumbling together at the base of the hydrozoans at night. Egg masses are usually deposited at the base of the hydrozoans, around each other.

**Life cycle.** Adults are present all over the year, especially in January–April and July–November; mating behavior and egg masses are recorded during January–April and June–December (based on 377 records, 116 sampling dates, and 28 different localities).

**Distribution.** South Africa (Shields, 2009); Belgium (OBIS, 2024); Italy (Schmekel & Portmann, 1982): Sardinia (Trainito & Doneddu, 2015); Greece (Thompson et al., 1990); Iberian Peninsula: Portugal, Andalucia, Levantine coast, Balearic Islands (Cervera et al., 2004); Catalonia (Ballesteros et al., 2016).

**Remarks.** This species exhibits a wide range of chromatic variability, the dorsolateral appendages’ background can be red, white, pink, or grey. *Doto rosea* is very similar to *D. moravesa* Ortea, 1997, *D. escatllari* Ortea, Moro & Espinosa, 1998, and *D. fragaria*. *Doto rosea* differs from *D. moravesa* for lacking black spots on tubercles and having black pigments at the base of some dorsolateral appendages (Ortea et al., 1997). As for *D. escatllari*, it has blue tones at the tips of the tubercles and thick spots on the back, whereas *D. rosea* has white tubercles and a body pigmented with fine dark dots (Ortea et al., 1997). The differences between *D. rosea* and *D. fragaria* are already mentioned in the section about *D. fragaria*. Given the morphological similarity of these four species, conducting a comparative molecular analysis would serve to reinforce their validity. Here, according to the phylogenetic and SDT results, *D. fragaria* and *D. rosea* are confirmed to be two different species.

***Doto verdicioi* Ortea & Urgorri, 1978**

(Fig. 18e–f)

*Doto verdicioi* Ortea & Urgorri, 1978: 79.

**Material examined.** SPAIN · 1 spc., L = 2 mm (sequenced); Galicia, Viveiro, Playa de Area; 43°41’36.2“N, 7°34’34.9”W, 2 m depth; 01 Jul. 2022; X. Salvador leg.; MCZ:Mala:406092.

**External morphology.** (Fig. 18e–f) Body elongated, narrow, translucent, white background, with brown irregular spots forming a longitudinal stripe on dorsum. Oral veil white pigmented on laterals. Rhinophores white pigmented on tip; rhinophoral sheath short, white pigmented on edge, base pigmented with irregular dots in internal part. Dorsolateral appendages arranged in 4–5 pairs; tubercles displayed in 3–4 crowns, short, almost smooth, with brown irregular dots scattered, except for apical white tubercle.

**Egg mass.** (Fig. 18f) White, displayed in a meandering cord shape. Depending on the substrate it can be either linear, with up to 4 folds, or deposited in a circle shape.

**Ecology.** The specimens were found on rocky substrates and over laminar algae with coverage of *Aglaophenia* sp., during the summer season.

**Distribution.** Cantabrian Sea, Galician coast, and Portugal (Cervera et al., 2004).

**Remarks.** In the original description, the rhinophores were described as having reddish spots and some white pigment at the apex (Ortea & Urgorri, 1978). In our specimens, this feature is not very noticeable. However, the presence of scattered brown spots on dorsolateral appendages can be observed well, except for the apical tubercle, which is white. These are diagnostic characters that easily distinguish *D. verdicioi* from other congeners.

***Doto vrenifossorum* sp. nov. Vázquez-Alcaide, Schrödl & Moles, 2025**

(Fig. 19)

**Fig. 19.**
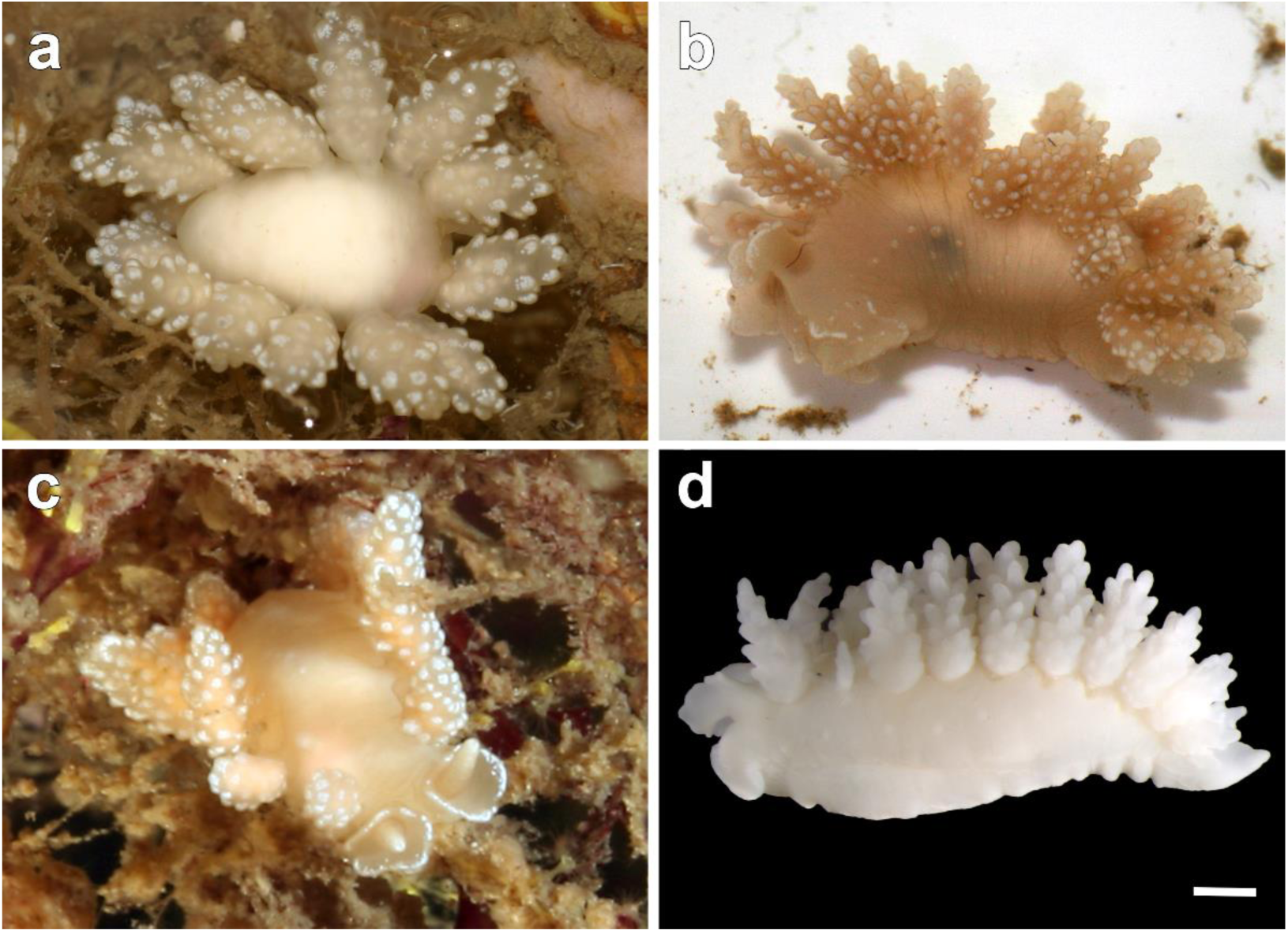
Photographs of specimens of the new species *Doto vrenifossorum* sp. nov. **a** specimen with whitish colouration **b** specimen with pinkish colouration **c** specimen found crawling on hydrozoans **d** specimen ZSMMol20130490 preserved. Scale bar: 1 mm. Photo (d) by Diego Vázquez-Alcaide, and photos of (a–c) courtesy of Vreni Häussermann & Fossi Försterra

**ZooBank registration link.** https://zoobank.org/NomenclaturalActs/771a810c-df80-472d-b6d3-dce5682f9306

*Doto* sp.: Schrödl, 2010: 534.

**Type material examined. *Holotype*.** SOUTH AMERICA · L = 5 mm (sequenced); Chile, Isla Royal Oak, West of Canal Messier; 47°58’45.0”S, 74°40’47.0”W, 27.5 m depth; 17 Mar. 2012; Günther Försterra leg.; ZSMMol20130494. ***Paratype*.** (sequenced); same collecting place and date as for holotype, 12.9 m depth; Günther Försterra leg.; ZSMMol20130495.

**Other material examined.** SOUTH AMERICA · 2 spcs, L = 10–18 mm (one sequenced and two dissected); Chile, Canal Copihue; 50°20’23.0”S, 75°22’39.0”W, 20 m depth; 16 Apr. 2013; Roland Meyer leg.; ZSMMol20130485 · 5 spcs, L = 7–10 mm (one sequenced and dissected); Chile, Muro Roberto; 20 m depth; 17 Apr. 2013; Roland Meyer leg.; ZSMMol20130490 · 1 spc., L = 13 mm (sequenced); Chile, Isla Zealous, East of Canal Messier; 47°56’54.01”S, 14°18’0.4”O, 13.2 m depth; 15 Mar. 2012; Günther Försterra leg.; ZSMMol20130496 · 1 spc. (sequenced); Chile, Canal Pitt; 50°50’49”S, 74°3’32”O, 5–20 m depth, 07 Mar. 2006; M. Schrödl & Roland Melzer leg.; ZSMMol20070371 · 1 spc. (sequenced); same collecting place and date as for preceding, 5–20 m depth; M. Schrödl & Roland Melzer leg.; ZSMMol20070369 ·1 spc. (sequenced); same collecting place and date as for preceding, 5–20 m depth; M. Schrödl & Roland Melzer leg.; ZSMMol20090486 · 1 spc., L = 7 mm (sequenced); Chilean Patagonia; CRBA-113028.

**Etymology.** This species is named after Vreni Häussermann and Fossi Försterra, eager explorers of the Chilean Fjord Region for almost three decades.

**Diagnosis.** Body background colour white to pinkish, with rows of small protuberances on the sides and head. Oral veil undivided and without projections. Rhinophores of same color the body with dull spot on tip.

Rhinophore sheath lobed with white dots on margin. Dorsolateral appendages elongate, distributed in up to 7–13 pairs; tubercles in 6–8 crowns, gnarled, with white pigmentation apically.

**External morphology.** (Fig. 19) Body elongated, bulky laterally, background colour white to pink; irregular rows of small knobs on body sides and head. Anal papilla large, located dorsally in mid-right position. Oral veil not divided, trapezoidal, without digitiform projections. Rhinophores digitiform, smooth, white to pink, with opaque white spot at tip. Rhinophoral sheath wide, smooth, lobate border with white dots on edge. Dorsolateral appendages distributed in up to 7–13 pairs, elongated, displayed in 6–8 crowns; tubercles knob-like, with white apical pigmentation. Small gill leaflets on inner side of dorsolateral appendages stems.

**Radula.** (Fig. 20) Radular formula 0, 1, 0; rachidian arched, pointed tip; between 5–7 denticles along margin, unequal among and within teeth, those closer to medial cusp pointed, those more distal blunt.

**Fig. 20.**
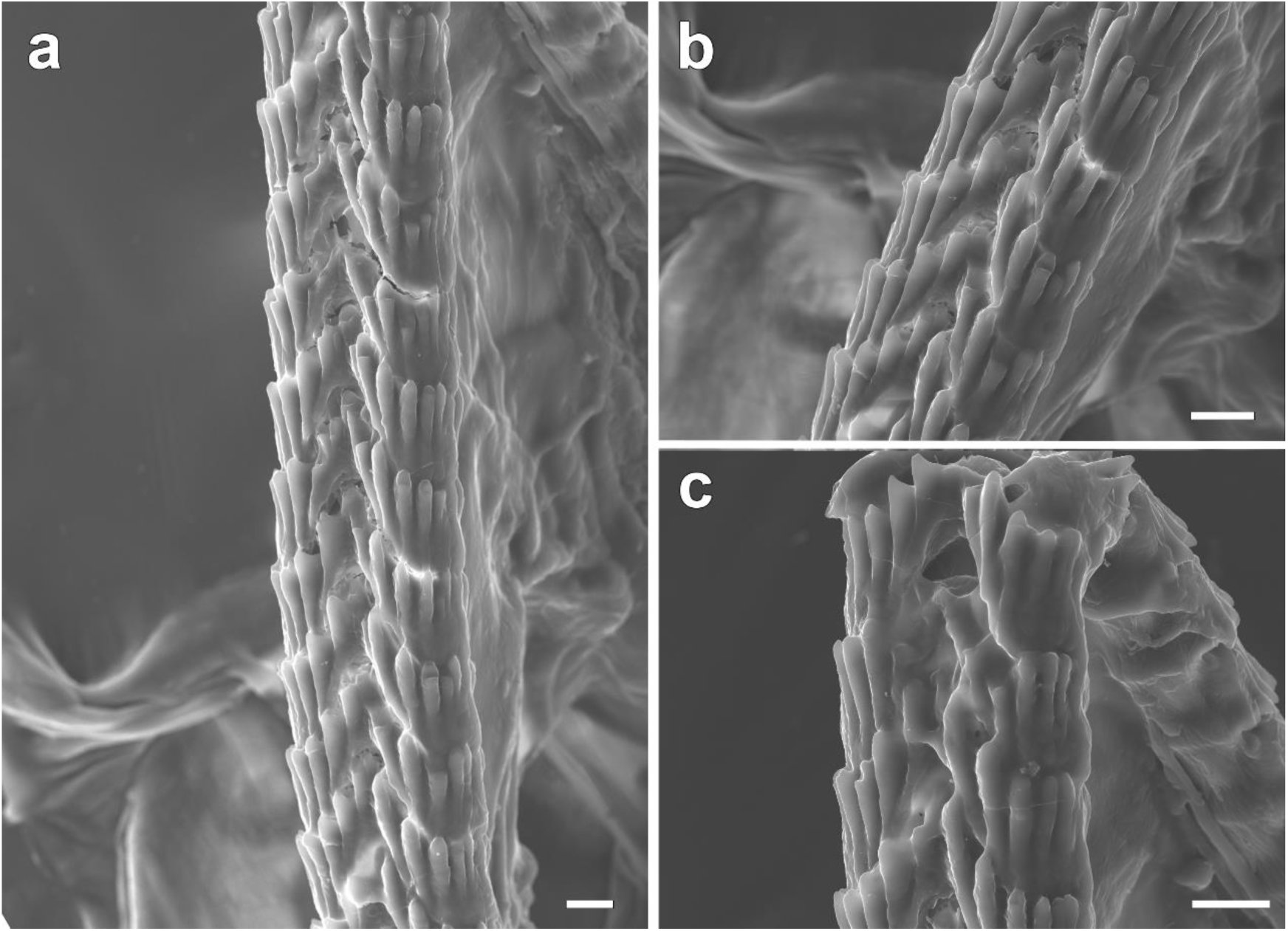
Scanning electron micrographs of the radula of *Doto vrenifossorum* sp. nov. (ZSMMol20130490) **a–c**. Scale bar: 10 μm. Photos by Diego Vázquez

**Reproductive system.** (Fig. 21) Ampulla bean-shaped below albumen gland. ♂ Prostate long, eigth times folded, attached to a slender, contorted vas deferens, connected to penis. Penis short, ovoid; penile sheath saccular. Genital openings situated in anterior-right position. ♀ Oviduct branching into ampulla, prostate and albumen gland (nidamental gland 1). Albumen gland (nidamental gland 1) large, globose, sharing a wide atrium with the vagina. Mucous gland (nidamental gland 2) globular in texture, above albumen gland, where prostate is located. Seminal receptacle saccular, elongated, attached to genital opening by a thin and long duct. Vagina large, broad, oval, reaching outwards through a wide stellate opening.

**Fig. 21.**
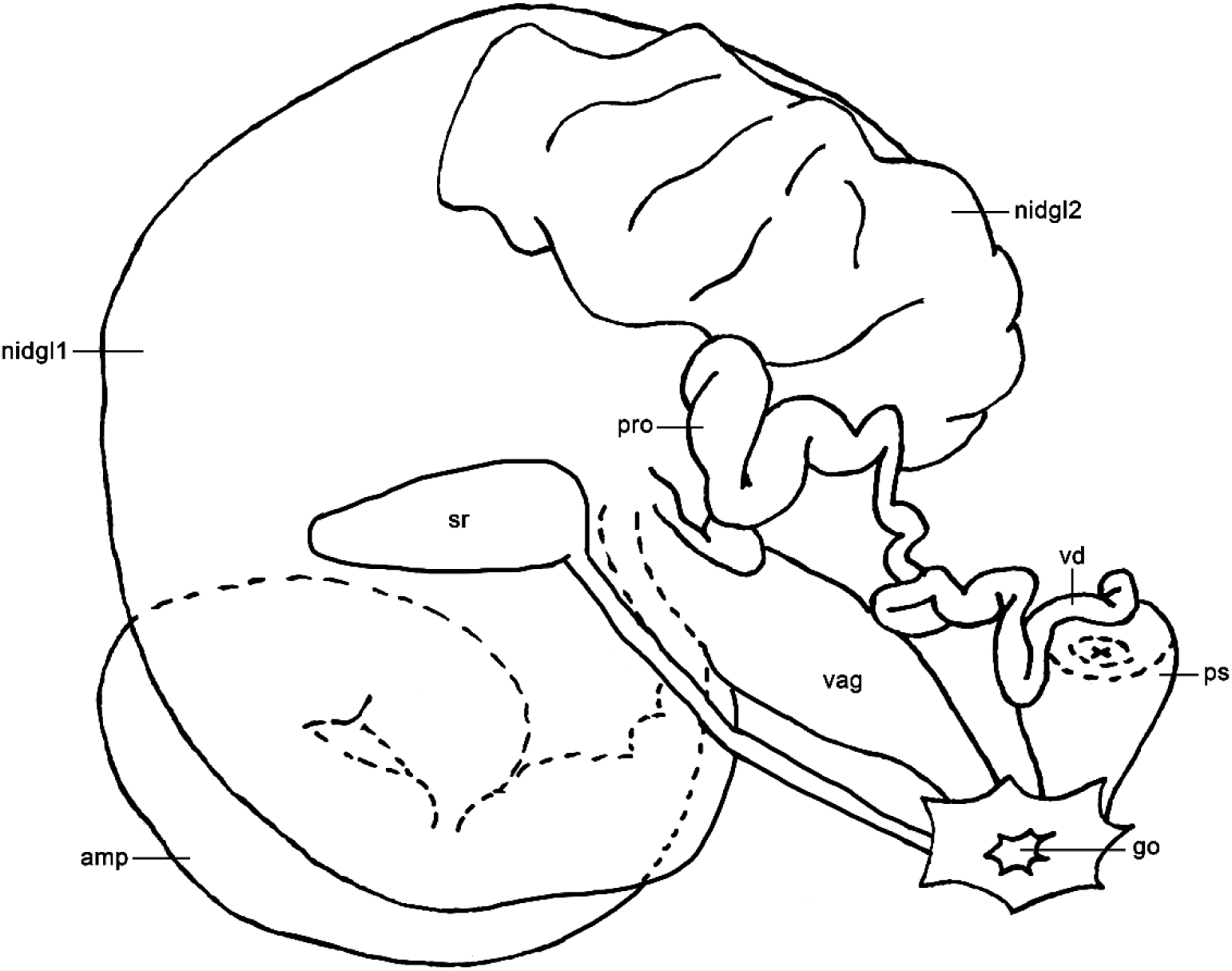
Schematic drawing of the reproductive system of *Doto vrenifossorum* sp. nov. Abbreviations: amp, ampulla; go, genital opening; nidgl1, albumen gland; nidgl2, mucous gland; ps, penial sheath; pro, prostate; sr, seminal receptacle; vag, vagina; vd, vas deferens

**Ecology.** The specimens were found on vertical rock walls with hydroids in different diving areas between 5–30 m depth, some specimens on steep walls exposed to strong currents.

**Distribution.** This species is only known from the Chilean Patagonia from the Aysén Province to the Southern Magallanes Fjord.

**Remarks.** Externally, *D. vrenifossorum* sp. nov. is very similar to *D. carinova* Moles, Avila & Wägele, 2016 from 277 m depth in the Weddell Sea, Antarctica (Moles et al., 2016). Besides the difference in distribution and depth, *D. vrenifossorum* sp. nov. has a lobed rhinophore sheath and protuberances on the sides and head, while *D. carinova* has a cylindrical rhinophore sheath and the protuberances on the body are absent (Moles et al., 2016). The reproductive system of *D. vrenifossorum* sp. nov. also resembles that of the Antarctic species *D. carinova*. However, they differ in that *D. vrenifossorum* sp. nov. has a short, ovoid penis, an elongated, saccular seminal receptacle, and a large, wide, oval vagina, whereas *D. carinova* has a long, conical penis, a small, rounded seminal receptacle and a short, flattened vagina (Moles et al., 2016).

## Discussion

The taxonomy of *Doto* species has been contentious due to their intraspecific morphotypic variability and interspecific similarity, which categorizes them as cryptic species (Shipman & Gosliner, 2015). In this study, we build upon the molecular data of *Doto* species found in the Mediterranean Sea, Eastern Atlantic, and South America. For these species, a comprehensive account of their external morphology, ecological characteristics, life cycle, distribution, and egg mass are provided. An integrative description of each species is of utmost importance, as *Doto* species have frequently been misidentified due to the absence of distinct diagnostic characters.

Sixteen *Doto* species have been documented in the Mediterranean (GROC, 2024; OPK-Opistobranquis, 2024), but only five have been tested with molecular data (Moles et al., 2016). Out of the 171 terminals included in our phylogeny, 59 are newly sequenced specimens belonging to 20 species. From these, 12 are found in the Mediterranean (*D. cervicenigra*, *D. coronata, D. eireana, D. floridicola, D. fragaria, D. koenneckeri, D. maculata, D. millbayana, D. paulinae, D. pygmaea*, and *D. rosea*), including the first new species described from the Catalan coast (*D. cavernicola* sp. nov.). Moreover, ten of the total number of species included have been sequenced here for the first time, from the Atlantic (*D. fluctifraga* and *D. verdicioi*), Mediterranean (*D. cavernicola* sp. nov., *D. cervicenigra*, *D. fragaria, D. pygmaea*, and *D. rosea*), Peru (*Doto* sp. 1 and *Doto* sp. 2) and the second new species described from Chile (*D. vrenifossorum* sp. nov.).

The records of the Mediterranean species included in this study are mainly based on the winter and spring months. This fact may be related to the life cycle of their hydrozoan prey. In the Mediterranean, the lowest temperatures in the water column occur between these months, and the highest in the summer (Brasseur et al., 1996). In general, Mediterranean hydrozoan species thrive in cooler waters, as they are sensitive to high temperatures, which limits their abundance (Puce et al., 2009). Consequently, during the summer months, hydrozoans enter a state of latency, leading to the end of the nudibranch life cycle.

### Phylogenetic relationships of the first offshoots, Clade 1, and Clade 2

Phylogenetic analyses reveal that *D. pinnatifida* clade forms a polytomy with the clades of the other *Doto* species, consistent with previous molecular studies (Shipman & Gosliner, 2015; Moles et al., 2016; Martinsson et al., 2021). Based on the histone H3 marker, *D. cuspidata* from the North Atlantic Ocean was recovered as the sister species of *D. pinnatifida* (Martinsson et al., 2021). We suggest that further studies with additional molecular markers should be conducted to better test the relationship of *D. pinnatifida* and *D. cuspidata* to other *Doto* species and to assess *D. cuspidata* species’ validity. Within the same early polytomy is *D. vrenifossorum* sp. nov., found in the fjords of Patagonia and described here for the first time. The species was previously recognized but not officially described until now (Schrödl, 2010). The morphological similarity to *D. carinova* from the Weddell Sea in Antarctica is striking, but the presence in *D. vrenifossorum* sp. nov. of protuberances on the sides and head, and the presence of a lobate rhinophore sheath (see Fig. 19), together with differences in the reproductive system, and its bathymetric and geographic distribution, aids at describing it as a new species (see remarks section above).

Clade 1A comprises *Doto* species from Antarctica, the Atlantic, Mediterranean, and Indo-Pacific Oceans. For the Antarctic species *D. antarctica*, this study provides additional molecular evidence from specimens collected in the South Shetland Islands, supporting its circumpolar distribution, as suggested in previous studies (Moles et al., 2016). The Atlantic and Mediterranean species within the clade, namely *D. fluctifraga*, *D. cervicenigra*, and *D. pygmaea*, are characterized by their dark colouration (Fig. 7, Fig. 11d, Fig. 11f, and Fig. 17). The case of *D. fluctifraga* and *D. cervicenigra* is a clear example of crypsis, as despite the lack of discernible differences in external morphology (see systematic descriptions), genetic results reveal that they are distinct species. Distribution and diet are important characters to differentiate both species. *Doto fluctifraga* has an Atlantic distribution and feeds on *Pennaria disticha* (Ortea & Pérez, 1982; Ortea et al., 2010; this study), while *D. cervicenigra* has a Mediterranean distribution and feeds on *Obelia*, *Campanularia*, and *Sertularella mediterranea* (Ortea & Bouchet, 1989; Lombardo & Marletta, 2020; Salvador, Fernández-Vilert & Moles, 2022; this study). This showcases the relevance of integrative taxonomy to differentiate species of *Doto*. Another species with black colouration is *D. pygmaea*, which can be found drifting on floating debris. This species has been documented with an Amphiatlantic distribution, also including the Mediterranean (Schmekel & Portmann, 1982; Ortea et al., 1997; Sanvicente-Añorve et al., 2012). Here, we confirm its presence in the Mediterranean Sea through molecular data, while its Atlantic distribution has yet to be validated. Probably *D. pygmaea* has a cosmopolitan distribution, like the nudibranch *Fiona pinnata* (Eschscholtz, 1831) that also inhabits floating objects (Trickey et al., 2016), due to the dispersal characteristics of its habitat.

Clade 1B includes two Mediterranean and four Southern Pacific species sequenced here. The distinctiveness of the Mediterranean *D. fragaria* and *D. rosea* has been validated with molecular data. While both species share similar external morphological characteristics, distinguishing them requires examination of the tubercle colouration. While *D. fragaria* has white tubercles with a black inner centre (Fig. 12 and Fig. 8a), *D. rosea* exhibits entirely-white tubercles (Fig. 18a–d and Fig. 8b). Additionally, the shape and colour of egg masses can aid in species identification. The egg mass of *D. fragaria* is white, short, and saccular, while that of *D. rosea* is longer, sinuose, and sometimes circular. Concerning the Pacific species, sequences of four unidentified or undescribed species are included, namely *Doto* sp. 1, sp. 2, sp. 3, and sp. 4. The latter groups with other unidentified specimens sequenced in previous studies, encompasses a wide distribution in the eastern Pacific, from Washington State to Peru, yet with unknown species identity. Overall, the presence of multiple undescribed species in the Western Pacific highlights the need for further taxonomic efforts in that region.

In Clade 2, composed of species from the North Atlantic Ocean, results are comparable to those in the study by Martinsson and collaborators (2021). This clade includes the *D. fragilis*, *D. hystrix*, and *D. formosa* species complex, which we suggest needs revision with more molecular, ecological, and morphological data than currently available to delimit and validate these species.

### Phylogenetic relationships of Clade 3 and *D. coronata* species complex

The Northeast Atlantic species *D. verdicioi* was sequenced here for the first time and found in a clade with *D. maculata*, previously documented with a North Atlantic distribution (Lemche, 1976; Urgorri & Besteiro, 1986). Here, we extend *D. maculata* distribution to the Western Mediterranean Sea. COI, used for the SDTs analyses, shows intraspecific divergence for the Mediterranean *D. maculata* specimens (see Fig. S1), contrary to 16S and H3 (see Fig. S2 and Fig. S3), albeit with an uneven number of sequences each. Increasing the number of sequenced specimens across its revised distribution could clarify this putative case of cryptic speciation. In the same clade, ML confirms the species validity of *D. paulinae* and *D. koenneckeri*, a pattern not supported by BI, possibly due to the low variability of the nuclear marker used. PTP delimits *D. paulinae* and *D. koenneckeri* as distinct species, whereas ASAP clusters them within the same clade. The mitochondrial markers support the results obtained by PTP.. Nevertheless, morphological differences also support keeping both species. *Doto koenneckeri* has elongated tubercles of dorsolateral appendages with brown comma-shaped spots (Fig. 13 and Fig. 8e), whereas in *D. paulinae* they are completely brown, except for the apical one (Fig. 16 and Fig. 8d). In addition, *D. koenneckeri* has white pigmentation on the rhinophoral sheath and pigmented bands on the sides of the body, both characters absent in *D. paulinae*. Based on these results, we maintain both species as valid.

However, we recognize that future studies incorporating a greater number of markers are needed to better clarify the relationships between these two species.Clade 3B includes the *D. millbayana* and *D. dunnei* species complex from the North Atlantic Ocean, and the morphotypes of *D.* ‘*coronata*’ and *D.* ‘*dunnei*’ from the Mediterranean. For the Atlantic specimens, the study by Shipman & Gosliner (2015) found no genetic differences to suggest that *D. millbayana* and *D. dunnei* were different species, recommending a larger taxon sampling to clarify their status. Later, Martinsson and collaborators (2021) obtained similar results by increasing the number of sequenced specimens from the Atlantic. The latter study considered the differences in diet, pseudobranch shape, body pigmentation, and radula enough to keep both species. In this study, we also increased the number of Mediterranean specimens and sequenced markers of *D. dunnei* morphotypes and *D. millbayana*. The phylogenetic tree and all SDT analyses show all specimens of the *D millbayana*/*D. dunnei* complex, along with their respective morphotypes mentioned, constitute the same species. Therefore, our molecular results and those of previous studies (Shipman & Gosliner, 2015; Martinsson et al., 2021), considering that *D. millbayana* was the first described species, support *D. dunnei* syn. nov. of *D. millbayana* (**new synonymy**). Molecular evidence also shows that the specimens widely and erroneously identified as *D. coronata* and *D. dunnei* from the Mediterranean correspond to *D. millbayana*. This set of results reports the wide morphological and chromatic variation of *D. millbayana* from the North Atlantic to the Mediterranean. The morphological variation of the species ranges from having dorsolateral appendages with rounded tubercles (traditionally attributed to *D. dunnei* and *D. millbayana*; see Fig. 15b–d and Fig. 5c2–3) to smoother dorsolateral appendages with less conspicuous tubercles (traditionally attributed to the Mediterranean *D. coronata*; see Fig. 15a, e–f and Fig. 5c2). On the other hand, the chromatic variation of the irregular markings ranges from purple to reddish, and these may be less frequent and more scattered (traditionally attributed to *D. millbayana* and the Mediterranean *D. coronata*) or numerous and closer together, and may even merge, causing some specimens to be almost completely pigmented (traditionally attributed to *D. dunnei*). Similarly, the tubercles of dorsolateral appendages may present one or more dots per tubercle (traditionally attributed to *D. millbayana* and the Mediterranean *D. coronata*) or one dot with discontinuous dotted circles around it (traditionally attributed to *D. dunnei*). There is also evidence that *D. millbayana* has a generalist diet, as is the case with *D. coronata* (Lemche, 1976; Picton & Brown, 1981; Shipman & Gosliner, 2015; Martinsson et al., 2021). Their diet includes *Aglaophenia* sp. (Martinsson et al., 2021), *Nemertesia ramosa* (Lamarck, 1816) (Thompson & Brown, 1984), *Plumularia setacea* (Linnaeus, 1758) (Lemche, 1976), *Sertularia argentea* Linnaeus, 1758 (Shipman & Gosliner, 2015), *Kirchenpaueria pinnata* (Lemche, 1976), *Tridentata perpusilla*, *Kirchenpaueria halecioides*, and *Obelia* sp. (this study).

Nevertheless, a single specimen of *D. coronata* was confirmed in the Mediterranean with molecular data, together with *D. eireana* and *Doto cavernicola* sp. nov. (clade 3B). Mediterranean specimens wrongly identified as *D. coronata* did not cluster with the specimens from the type locality in the North Atlantic (Moles et al., 2016). Similarly, most of the specimens identified as *D. coronata* in the Mediterranean correspond to *D. millbayana*, except for one isolated specimen found living in cave entrances, in sympatry with *D. cavernicola* sp. nov. Therefore, our molecular results and those of previous studies confirm the distribution of *D. coronata* on both sides of the North Atlantic (Verrill & Smith, 1874; Shipman & Gosliner, 2015; Thompson et al., 1990; Ortea et al., 2008) and the Mediterranean (this study), and reveal that its Mediterranean distribution is more restricted than previously thought. The specific Mediterranean specimen of *D. coronata* has a high morphological similarity to *D. cavernicola* sp. nov. (see remarks section). However, the phylogenetic tree and the SDT analyses in this study, together with differences in the reproductive system, distinguish both species.

The possibility of this singleton being a contamination can be ruled out since this belongs to the same clade as the previously sequenced specimens from the type locality, in the Netherlands (Shipman & Gosliner, 2015), and not sequenced here. Furthermore, the specimens of *D. cavernicola* sp. nov. are closely related to those of *D. eireana* and not to those of *D. coronata*. Also, *D. cavernicola* sp. nov. feeds specifically on *Diphasia* cf. *delagei*, while *D. coronata* has a more generalist diet (Lemche, 1976; Picton & Brown, 1981; Shipman & Gosliner, 2015; Martinsson et al., 2021). The diet specificity of *D. cavernicola* sp. nov. may be a diagnostic feature, as some *Doto* species are specialised to feed on a hydrozoan species, such as *D. maculata* which feeds specifically on *Halopteris catharina* (Johnston, 1833) (Lemche, 1976; Picton & Brown, 1981; Thompson & Brown, 1984; Martinsson et al., 2021). Concerning *D. eireana*, this species is documented to have a North Atlantic and Mediterranean distribution (Lemche, 1976; Ortea & Urgorri, 1978; Calado et al., 2003; Ballesteros et al., 2016), but so far only Atlantic specimens from NE Spain have been confirmed by molecular data (Wollscheid-Lengeling et al., 2001; Shipman & Gosliner, 2015; this study). Nevertheless, we suggest that its distribution in the Mediterranean may be based on misidentifications and likely corresponds to *D. maculata*.

Specimens of *D. eireana* are also morphologically very similar to *D. coronata* and *D. cavernicola* sp. nov. The former is distinguished by the fact that sometimes lacks apical dots on the apical tubercle of the dorsolateral appendages, these being wider below the tip (see Fig. 5d and Fig. 10). Additionally, *D. eireana* and *D. cavernicola* sp. nov. do not overlap in their distribution, while the former is only verified in the Atlantic with molecular data, *D. cavernicola* sp. nov. has only been found in the Mediterranean.

In summary, our results underscore the complexity of *Doto* taxonomy by highlighting substantial intraspecific morphological variability and cryptic diversity across the Mediterranean, Eastern Atlantic, and South American regions. Integrating molecular, morphological, ecological, and reproductive data has proven crucial for accurately delineating species boundaries, describing new species (e.g., *D. cavernicola* sp. nov. and *D. vrenifossorum* sp. nov.), and revealing misidentifications stemming from incomplete diagnostic characters. We also propose new synonyms (e.g., *D. dunnei* syn. nov. of *D. millbayana*) and extend distributional ranges for several species, clarifying previously uncertain records. Nevertheless, a wider taxon sampling—including understudied regions worldwide—remains essential to unravel the full scope of *Doto* diversity. Overall, these findings emphasise the need for continued integrative approaches to resolving the genus’s intricate taxonomy, especially given the broad morphological plasticity, overlapping colour patterns, and shared ecological niches that often mask true biodiversity.

## Contributions

The study was conceived by JM. Samples were provided by XS, MS, JM, and YH. Photographs were provided by XS. Lab work was conducted by JM and DV-A. Figures were designed by DV-A and XS. Literature research was conducted by DV-A, JM, and XS. Data analysis, interpretation, and writing of the first draft were done by JM, XS, and DV-A. Critical review of the article was carried out by JM, XS, GG, and MS. Funds were secured by JM, GG, and MS. All authors read and approved the final manuscript.

## Acknowledgements

We want to thank the citizen collaborators of GROC who regularly monitor the Catalan coast. We also thank R. Fernández-Vilert, G. Mas, I. Figueroa, G. Försterra, R. Meyer, R. Melzer, and J. Fernàndez for providing samples. Special thanks are due to J. Fernández-Simón, A. Enguídanos, and the members of Slug Lab (@slug_lab) for their help in the molecular lab and to the CCiT-UB for helping during the SEM sessions. We acknowledge the Catalan and Canary Islands local Governments for providing collection permits SF/0589/2018 and DG051201-333/2022, and SGBTM/BDM/AUTSPP/13/2023, respectively.

## Ethics approval

No approval of research ethics committees was required to accomplish the goals of this study because experimental work was conducted with an unregulated invertebrate species.

